# Structural determinants within the adenovirus early region 1A protein spacer region necessary for tumourigenesis

**DOI:** 10.1101/2020.06.23.168369

**Authors:** David P Molloy, Roger J Grand

## Abstract

It has long been established that group-A adenoviruses (HAdV-A12, -A18 and -A31) can cause tumours in new-born rodents with tumourigenicity related to the presence of a unique spacer region located between conserved regions 2 and 3 within the Ad12E1A protein. Group B adenoviruses are weakly oncogenic whereas most of the remaining human adenoviruses are non-oncogenic. In order to understand better the relationship between the structure of the AdE1A spacer region and oncogenicity of HAdVs the structure of synthetic peptides identical or very similar to the adenovirus12 E1A spacer region have been determined to be α-helical using NMR spectroscopy. This contrasts significantly with some previous suggestions that this region is unstructured. Using available predictive algorithms, the structures of spacer regions from other E1As were also examined and the extent of predicted α-helix was found to correlate reasonably well with the tumorigenicity of the respective viruses.

**Importance:** This research analysed small peptides equivalent to a region within the human adenovirus early region 1A protein that confers, in part, tumour inducing properties to varying degrees on several viral strains in rats and mice. The oncogenic spacer region is alpha-helical, which contrasts with previous suggestions that this region is unstructured. The helix is characterised by a stretch of amino acids rich in alanine residues that are organised into a hydrophobic or ‘water-hating’ surface that is considered to form a major site of interaction with cellular protein targets that mediate tumour formation. The extent of alpha-helix in E1A from other adenovirus species can be correlated to a limited degree to the tumourigenicity of that virus. Some serotypes show significant differences in predicted structural propensity suggesting the amino acid type and physicochemical properties are also of importance.

## Introduction

Human adenoviruses (HAdVs) are dsDNA, non-enveloped viruses curated into 7 clades (A-G; Table 1) of 90 genotypes (1). In general, HAdVs -A12, 18, 31 and presumably, - A61 (2), can induce malignant tumours 2-5 month’s post-infection in neonate hamsters (3, 4) and mice (5–7). Serotypes -B3, 7 and 11 result in a lower frequency of tumours over prolonged periods (*ca* ~1 year; 8, 9). Group-C, -E and -F viruses are considered non-oncogenic, as are most HAdV-Ds, except for –D2:9 and:10 which induce post-infection development of oestrogen-dependent mammary tumours in female rats (10–12). Notwithstanding, serotype -G52 is considered non-oncogenic (Table 1; 13).

**Table 1.** Adenovirus subgroups

| Group | Adenovirus serotype | Relative Oncogenicity |
| --- | --- | --- |
| A | 12, 18, 31 | Strong |
| B:1 | 3, 7, 16, 21, 50 | Weak |
| B:2 | 11, 14, 34, 35, 55 | Weak |
| C | 1, 2, 5, 6, 57 | None |
| D:1 | 25, 60, 30, 33 | None |
|  | 39, 59, 19 |  |
|  | 8, 54 |  |
|  | 49 |  |
|  | 64, 37, 53 |  |
|  | 46, 24, 22, 20, 69, 42 |  |
|  | 43, 48, 58 |  |
| D:2 | 10, 9 | (Weak) |
|  | 27, 26, 29, 56 | None |
|  | 28, 32, 9 |  |
|  | 13, 47, 36 |  |
|  | 44, 45 |  |
|  | 38, 23, 51 |  |
| E | 4 | None |
| F | 40,41 | None |
| G | 52 | Not determined |

Most HAdVs and Ad DNA can be used to transform rodent cells *in vitro*, establishing distinct cell lines. Immunocompetent cells transformed by oncogenic serotypes can display tumourigenesis, although those transformed by non-oncogenic HAdVs remain so (reviewed 14–16). Most Ad-transformed cell lines will generate post-injection tumours *in vivo* in either immunocompromised nude mice, or syngeneic hosts (reviewed 14–16). Consequently, HAdVs are deemed to be oncogenic in rodents, with the viral Early Region 1A (E1A) gene product implicated in tumour development within susceptible hosts (reviewed 14–17). AdE1A expression alone can transform rodent cells *in vitro* although, transformation efficiency is markedly increased in cooperation with a complementary oncogene including, AdE1B or activated ras (18–23). Despite this, several Ad-transformed human cell lines exist where AdE1A exerts an opposing influence of tumour suppression (24).

AdE1A is the first protein expressed during viral infection and presents an intrinsically disordered multiplex hub to coordinate viral protein synthesis, cell cycle progression and facilitate cellular transformation (reviewed 25–27). Important interactions with cellular regulators are mediated through short linear motifs (SLiMs; 28) dispersed across 5 domain and 3 inter-domain regions (IDRs) present within most E1As (Fig. 1A; reviewed 29). Target proteins total some 57 to date and include the S4 proteasome subunit and nuclear GTPase, Ran which, interact exclusively at the N Domain (Fig. 1A; 29). Other interactive proteins including the cAMP response element-binding protein, CBP and related p300, display dual specificity in binding to both the N Domain and conserved region 1 (CR1, Fig. 1; reviewed 29). Similarly, formation of the E1A•p105(Rb-pocket)•p300TAZ-2 ternary complex (30) involves simultaneous binding interactions with motifs present within CR1 and CR2 (25–27, 30).

**Figure 1:**
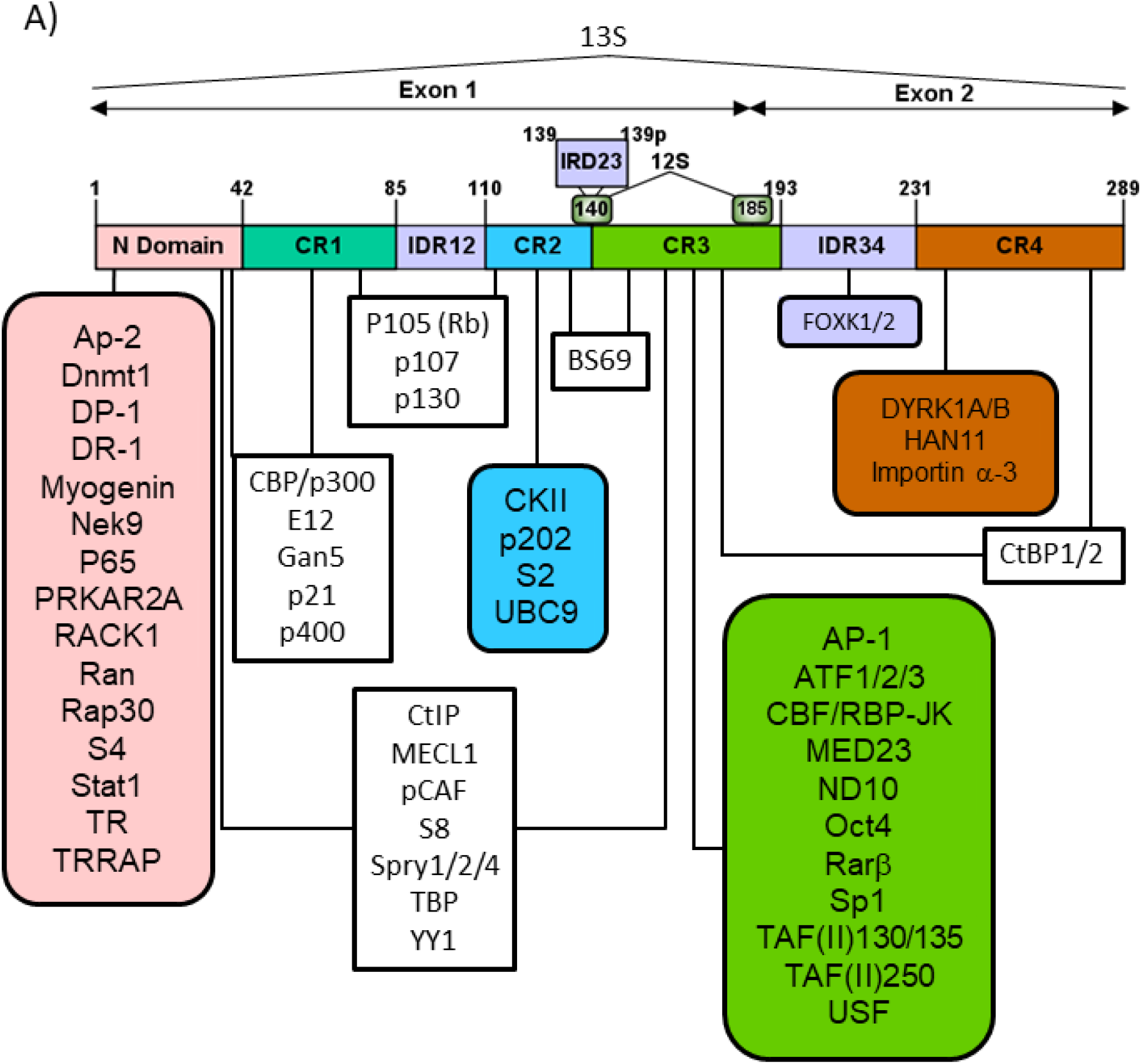
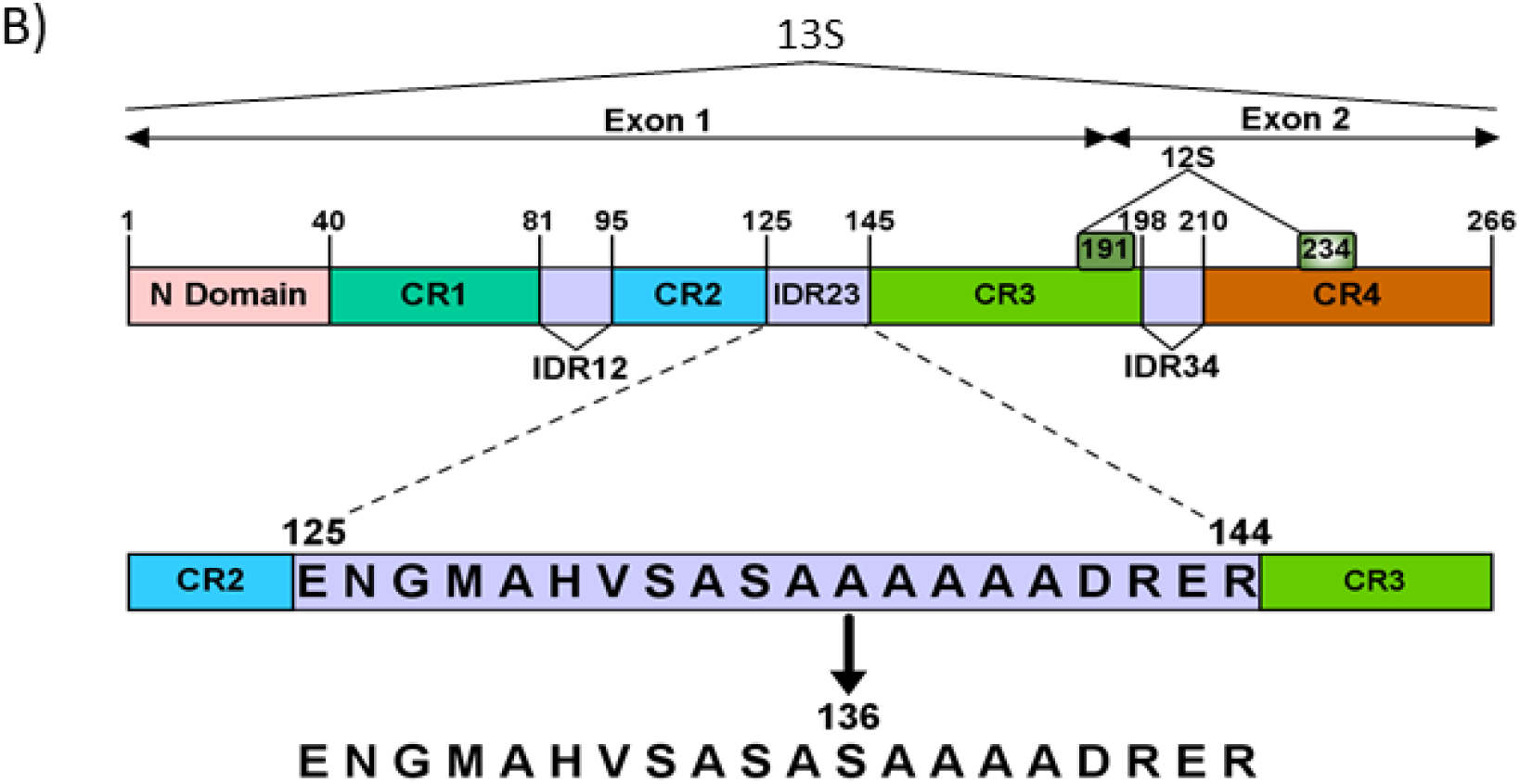
The structure of adenovirus E1A. A) The domain architecture of Ad5 13S E1A is represented as a linear cartoon sequence of 289 a.a.’s encoded within 2 exons illustrated as filled blocks for N-Domain (pink) and conserved regions (CR) 1 (dark green), 2 (blue), 3 (light green) and 4 (brown) along with inter-domain regions, IDR12, IDR23 and IDR34 (light purple). Domain boundaries are numbered at the N terminal sequence position for each above the primary sequence. Alternate splicing generates the smaller 12S product illustrated by the absence of residues between 140 and 185 (˄). The approximate location of target proteins within each domain are shown below the primary sequence illustrated as either specific for a particular domain (filled blocks) or multiple binding locations (unfilled blocks). B) The structure for the 266a.a. Ad12 E1A protein is illustrated as for (A). The sequences of two 20a.a. synthetic peptides equivalent to wild type (upper sequence) and A136S (lower sequence) IDR23 from Ad12 E1A used for NMR spectroscopy in this study are presented below the below cartoon using the single letter abbreviation.

The CR3 transcriptional activation domain of E1A is required for efficient expression of other viral early region genes through interactions with cellular transcription factors (TAFs) and TBP (Fig. 1A; 31–33). This region is only present in its entirety in Ad5E1A when translated from a 13SRNA transcript (25) although, a second major alternate splice product is also expressed from a 12S message (Fig. 1; 17). A further site of significance located very close to the C-terminus of the protein, Conserved Region 4 (CR4) which provides sites of interaction with, amongst others, DYRK1A kinase (34) and HAN11 (Fig. 1; 35). In concert with structural elements present in CR3 (36), CR4 also forms a major binding site for the cellular co-repressor CtBP (37, 38) and contains a nuclear localisation signal (NLS; Fig. 1A).

Rigorous amino acid sequence analyses (29) have allowed redefinition of the ‘spacer’ region between CR2 and CR3, originally linked to oncogenicity (39–41) and virtually absent in Group-C E1As, as inter-domain region 23 (IDR23, Fig. 1; 29). Similar regions of homology are present in the oncogenic simian virus SAdV7 and other members of Group-A adenoviruses (32, 33). Amino acid sequence conservation of Group-A E1As is limited in the weakly-oncogenic HAdV-B:1/-B:2 and non- or weakly-oncogenic Group-D E1As (Table 1; 32, 33). In essence, there appears to be a direct correlation between the presence and/or integrity of IDR23 and viral-induced tumourigenesis in rodents, although, the Ad12 spacer by itself is not sufficient to induce tumours (40, 42). It has also been demonstrated that inclusion of the Ad12 spacer in Ad5/Ad12 chimeric E1A does not induce tumours (40, 42) as this also relates to species-specific N domain sequences (40, 43). Notwithstanding, higher levels of cytotoxicity and replication efficiency observed with HAdV-C5 may also favour cell death over tumour production (43).

In order to understand of the role of IDR23 in oncogenesis, the Ad12E1A sequence was examined by NMR spectroscopy. Structure predictions for E1A-IDR23 from various serotypes were also generated and used alongside bioinformatics protocols to mimic the physicochemical properties of amino acids that might contribute to AdE1A oncogenicity, and by extrapolation, different adenovirus species.

## Materials and Methods

### Synthetic Peptides

Two synthetic peptides identical or similar in sequence to the spacer region of wild type Ad12E1A used in this study (Figure 1) were synthesised using standard f-moc procedures and purified by high performance liquid chromatography on a Vydac C_18_ column eluted with a gradient of acetonitrile (0-100%) containing 0.1% (v/v) trifluoroacetic acid.

### Sequence alignments, secondary structure predictions and peptide modelling algorithms

Sequence alignments were prepared essentially as described (1, 29, 32, 33) with sequences for HAdV E1As obtained from http://ncbi.nlm.nih.gov. The alignments for a total of 70 E1A proteins across all serotype groups were derived by comparisons between the MUSCLE and Clustal Omega (https://www.ebi.ac.uk; 44) algorithms with MAFFT (https://mafft.cbrc.jp/; 45), set to default parameters and constrained to the input reference sequence for Ad5 289a.a. E1A. Protein logos were computed for representative E1A-IDR23s across all HAdV serotypes and within separate subclade groups using Weblogo3 (http://weblogo.threeplusone.com/; 46) referenced to a full scale of 4.3 bits of information within the full-length Ad5 13S E1A protein as described (29). IUPRED values, isoelectric points (pH(I) and grand average of hydrophobicity (GRAVY) scores for E1A-IDR23 amino acid sequences were calculated using the ProtPARAM web server (https://web.expasy.org/protparam/.; 47). Structure predictions for IDR23s were derived for representative E1A-IDR23s from each adenovirus serotype using PSI-PRED (bioinf.cs.ucl.ac.uk/psipred; 48), Jpred4 (www.compbio.dundee.ac.uk/jpred; 49), and PREDATOR (https://npsa-prabi.ibcp.fr; 50) fold recognition algorithms. From these, weighted averages for each amino acid within every sequence were generated.

For the Ad12E1A IDR23 mutant peptide A138P, IntFOLD, QUARK, and I-TASSER (zhanglab.ccmb.med.umich.edu; 51) programs were used to generate first approximation *ab initio* theoretical model structures. Goodness of fit and molecular dynamics of initial models were evaluated using GROMACS (52) to provide templates for refined rounds of calculations using weighted averaged structural constraints. The final 3-D model was predicted using Monte Carlo simulations and evaluated by Ramachandran analysis in PROCHECK (servicesn.mbi.ucla.edu/PROCHECK/) with > 96% of all residues in favoured/allowed molecular constraint regions. Protein structure analyses and visualisations were performed using UCSF Chimera (53) and swisspdbv (https://spdbv.vital-it.ch; 54).

### NMR spectroscopy

All NMR spectra were recorded on a Bruker DMX 500 MHz spectrometer. One-dimensional ^1^H-NMR spectra were recorded using a gated pre-saturation pulse of 1.5s duration (to remove the H_2_O signal) for accumulation over a 5500-Hz sweep width with a 90° pulse of 5μs. Spectra were recorded as free induction decays (between 1024 and 2048 transients) and Fourier transformed using a 2-Hz line broadening function, essentially as described previously (55–58).

Two-dimensional experiments were acquired with 2048 data points in F2 with a sweep width of 11ppm and with between 480-608 rows in F1. Solutions of peptides were 2mM in concentration routinely at 285K at pH5.5 in 50% (v/V) d_3_-TFE/40% H_2_O/10% ^2^H_2_O. The water resonance was suppressed by very weak pre-saturation applied through the WATERGATE sequence and solvent artifacts were suppressed by using pulsed field gradients. Total correlation spectroscopy (TOCSY) used an MLEV-17 mixing pulse of 60ms duration (10 KHz spin locking field). Nuclear Overhauser effect spectroscopy (NOESY) experiments were performed using a mixing time of 200ms duration. Between 128 and 256 transients were collected for TOCSY and NOESY experiments. Proton signals in TOCSY experiments were assigned to identify spin systems within individual residues, which were coupled to sequential (d_αN_ i, i+1) NOE cross-peaks in NOESY spectra. The volume of each cross-peak was integrated to estimate the distances between individual proton signals within NOESY experiments. The distances were grouped into five classes, strong (1.8-2.5Å), strong-medium (1.8-3.5Å), medium (1.8-4.5Å), weak (1.8-5.5Å) and very weak (1.8-6.5Å), with strong and medium NOEs allocated the same distance constraints in structural calculations.

The movement of chemical shifts of backbone amide protons with changes in temperature between 285 and 305K in TOCSY experiments were used to assess backbone hydrogen bonding patterns within the peptides and were linear in every case. ^3^J_HNα_ coupling constants were calculated from two-dimensional COSY experiments.

The structures of peptides were determined using X-PLOR NIH 3.1 and 3.8 available at https://nmr.cit.nih.gov/xplor-nih essentially as described (59, 60). Initial structures were computed using the *dg_sub_embed* subroutine, in which the CA, HA, N, HN, CB* and CG* atoms were embedded. The remaining atoms were then placed by template fitting and the atomic co-ordinates were allowed to evolve under the applied NOE distance constraints during the *dgsa* and *refine* simulated annealing subroutines (60). From 70 structures generated, approximately 25-30 were discarded as these demonstrated wrong-handedness in the distance geometry routine. The remaining structures were chosen with no violations of the applied NOEs of greater than 0.5Å with a final 14 structures accepted in the *accept* subroutine.

## Results

E1As are generally considered to be disordered polypeptides with structural propensity limited to short linear motifs (29) and the CR3 transactivation domain (1, 29). Many studies have, however, successfully employed crystallographic techniques and NMR spectroscopy to analyse Ad5/12 E1A site-specific structures (30, 55–58). Recent bioinformatics approaches have also enabled precision mapping of E1A domain architecture (Fig. 1A; 29) and reclassification of Group-D viruses as two distinct subclades (Table 1; 1). Here, AdE1As were examined with special reference to the 3-D structure and physicochemical properties of the Ad12 E1A-IDR23 ‘oncogenic spacer’ that might provide insights into the structural basis of tumourigenesis (Fig. 1B).

### Structure of the 20-Residue Ad12 E1A IDR23 ‘spacer’ peptide

The structure of a 20-residue peptide identical to wild type Ad12 E1A-IDR23 amino acids E^125^—R^144^ (Fig. 1B) was examined by ^1^H NMR spectroscopy. TOCSY and NOESY (of mixing time 200-ms duration) spectra of the peptide are shown and NOEs observed are summarised in figure 2. Low values for ^3^J_NH-α_ coupling constants for residues over the region V^131^–D^141^ are indicative of regular secondary structure within the backbone conformation of the peptide (Fig. 2B(i)). In addition, low temperature coefficients (<5ppb/K) observed for shifts in backbone amide proton signals are indicative of a regular CO_*i*_–NH_*i*_, _*i*+ 4_ donor-acceptor intramolecular hydrogen-bonding pattern as found within an α-helix (Fig. 2B(ii)). Based on the types and relative volumes of NOE cross-peaks namely, *d*_αN_(*i*, *i* + 3) NOEs G^127^–H^130^, M^128^-V^131^, A^129^-S^132^, H^130^–A^133^, V^131^–S^134^, S^132^–A^135^, A^133^– A^136^, A^134^–A^137^, A^139^–R^142^ and *d*_αN_(*i*, *i* + 4) NOEs N^126^–H^130^, A^133^–A^137^ (Fig. 2B(iii)), region V^131^–D^141^ of the peptide conforms to a series of tight overlapping, right-handed α-helical turns.

**Figure 2:**
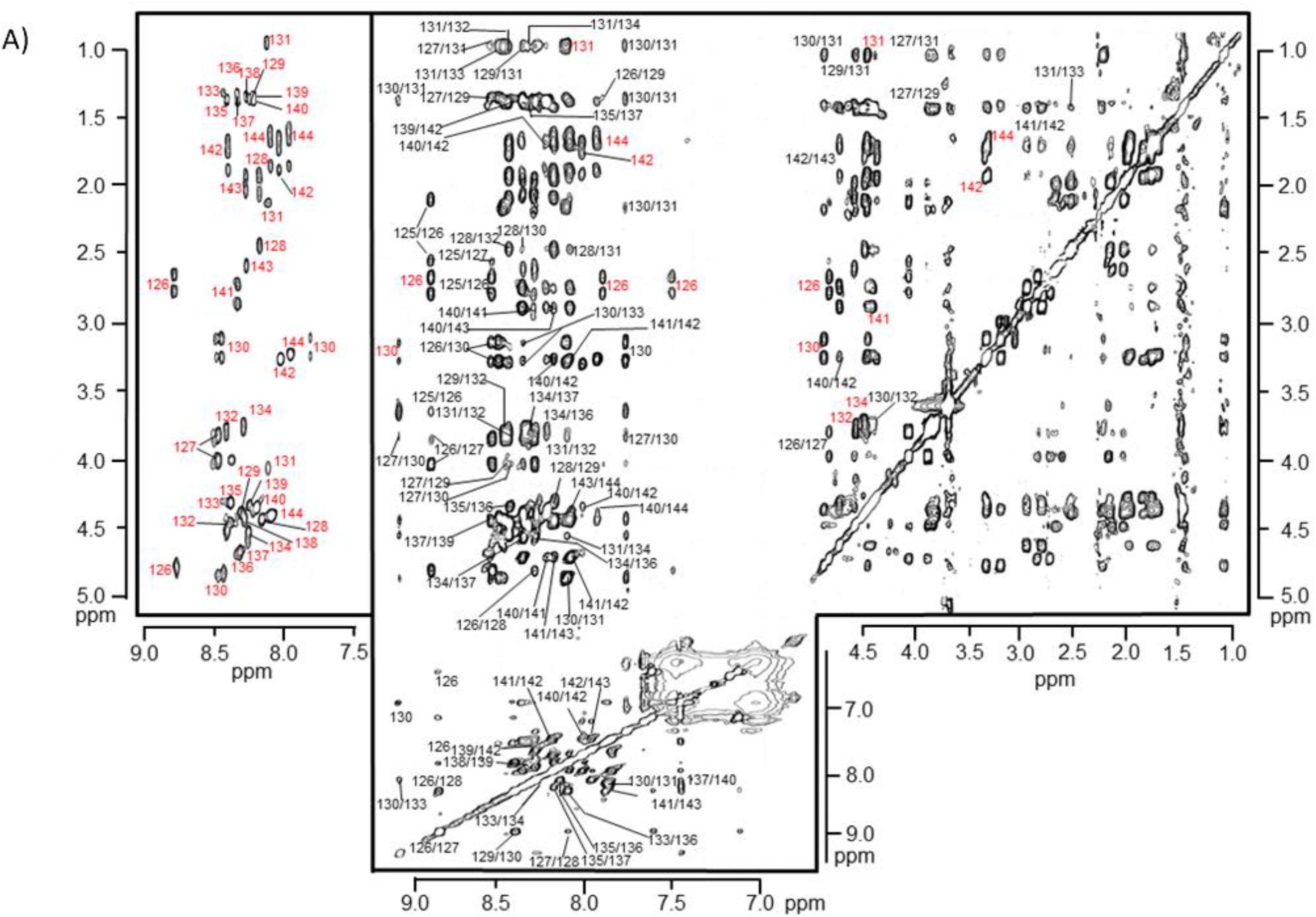
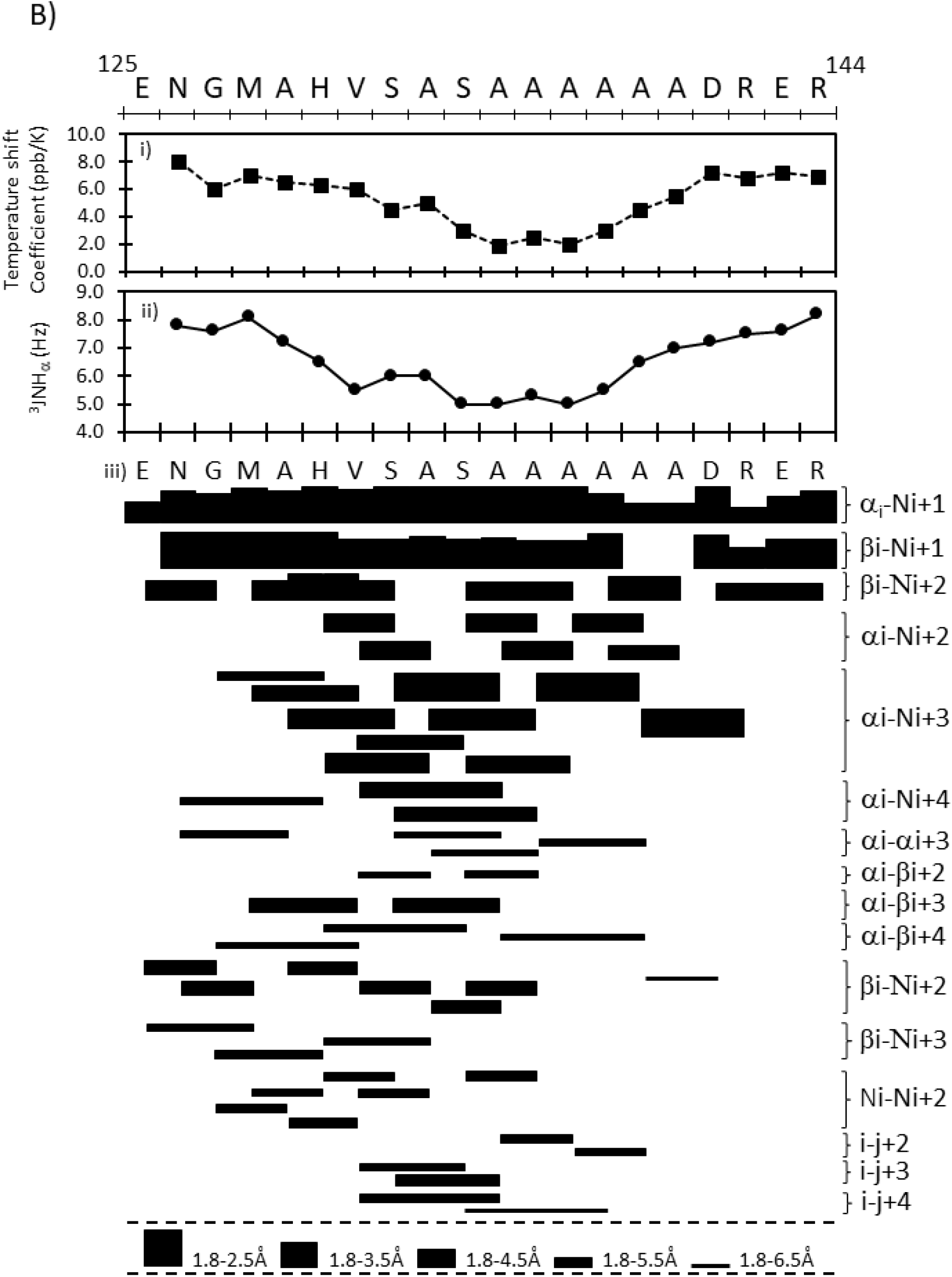
^1^H NMR assignments for the 20-a.a. wild type Ad12 peptide. A) In the far left hand panel the backbone amide-α region of a TOCSY experiment performed on the wild type form of the Ad12 13S E1A spacer region peptide is illustrated. Individual amino acids are labelled according to their type and position within the full-length Ad12 13S E1A protein; individual residues are labeled in red typescript. The NOESY spectrum (mixing time, 200 ms duration) for the wild-type peptide is presented in the large panel. The experiment was performed on a sample of the peptide at a concentration of 7 mg/ml in 50% (v/v) TFE/40% H_2_O/10% ^2^H_2_O, pH 5.5, at 285 K. NOEs corresponding to cross-peaks within a single residue are labeled with a single number corresponding to those positions in the sequence of full-length Ad12 E1A. NOEs that define medium range connectivities of the type d_αN_(i, i+2),d_βN_(i, i+2), and *d*_αN_(*i, i* +3) are labeled with both residue numbers; intra-residue signals are labelled in red typescript. B) Summary of the NOEs observed for the 20-residue wild type Ad12 spacer peptide. The temperature shift coefficients (i) and dihedral constraints (ii) are illustrated in graphical format above (iii) the distance (Å) between the sequential ((*i*, *i* + 1)), medium ((*i*, *i*+ 2) and (*i*, *i* + 3)), and long range ((*i*, *i* + 4)) NOEs observed are represented by the thickness of the bars. NOEs of the type (*i*, *j* + 2) (*i*, *j* + 3), and (*i*, *j* + 4) indicate classes other than *d* _αN_, *d* _NN_, or *d* _βN_.

Fourteen superposed structures from a conformational ensemble of the 40 lowest RMSDs calculated for the wild-type peptide reveal that the backbone conformation over the N- and C-terminal sequences (residues E^125^–M^128^ and R^142^–R^144^, respectively) of the peptide form frayed ends (Fig. 3A). However, between residues A^129^ and D^141^, a more ordered conformation in the peptide backbone is observed such that R-groups of amino acids involved in helix formation are unidirectional and conform the classical α-helix side-chain orientation (Fig. 3A) by virtue of overlapping turns formed within the peptide backbone (Fig. 3B). When considered in a clockwise N-to C-terminal helical wheel projection, the amphipathic nature of the helix displayed within the averaged calculated structure is exemplified over the region M^128^―D^141^ with alanine sidechains at positions 129, 133, 136, 137 and 140 forming a distinctive hydrophobic surface (Fig. 3C).

**Figure 3.**
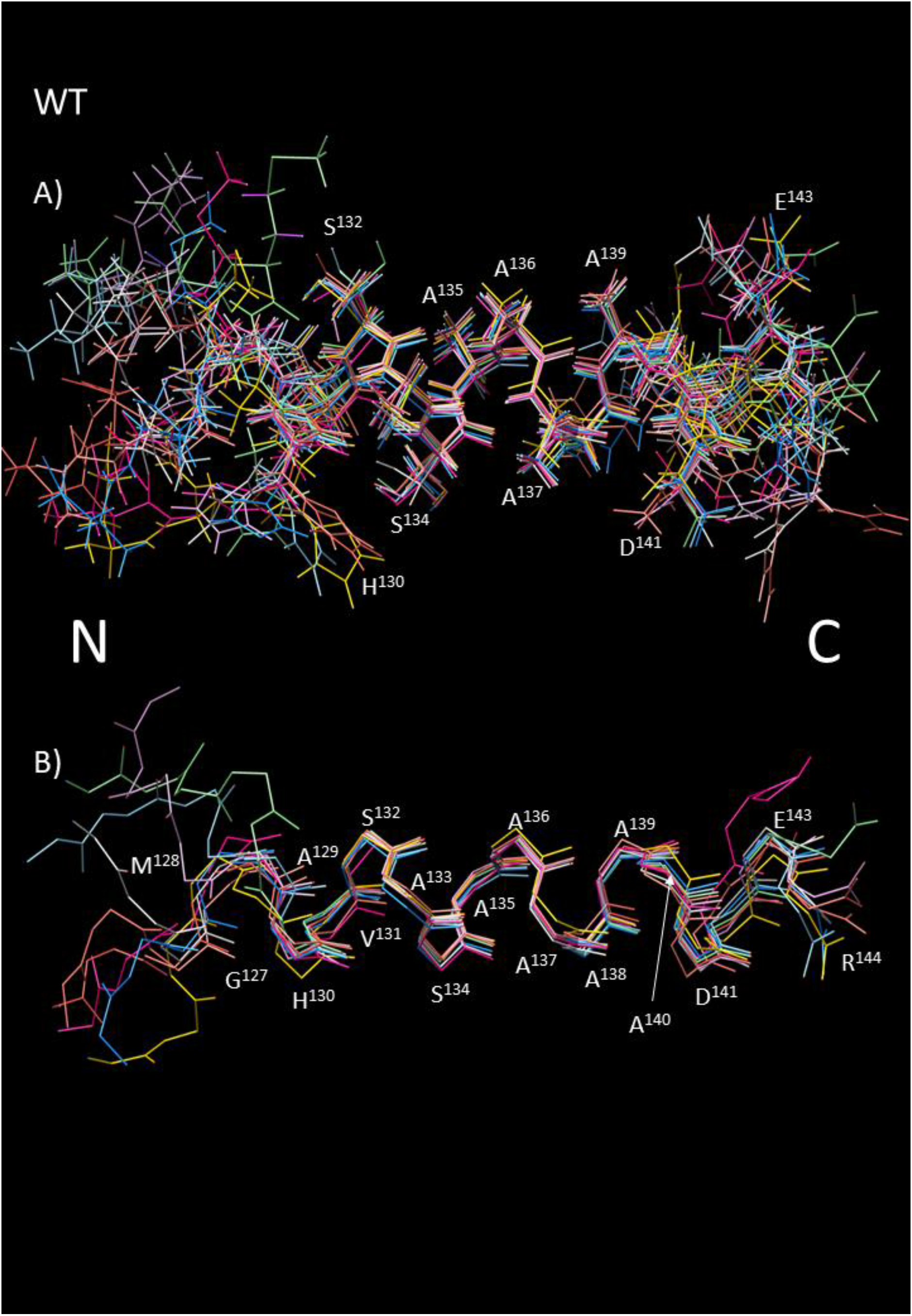
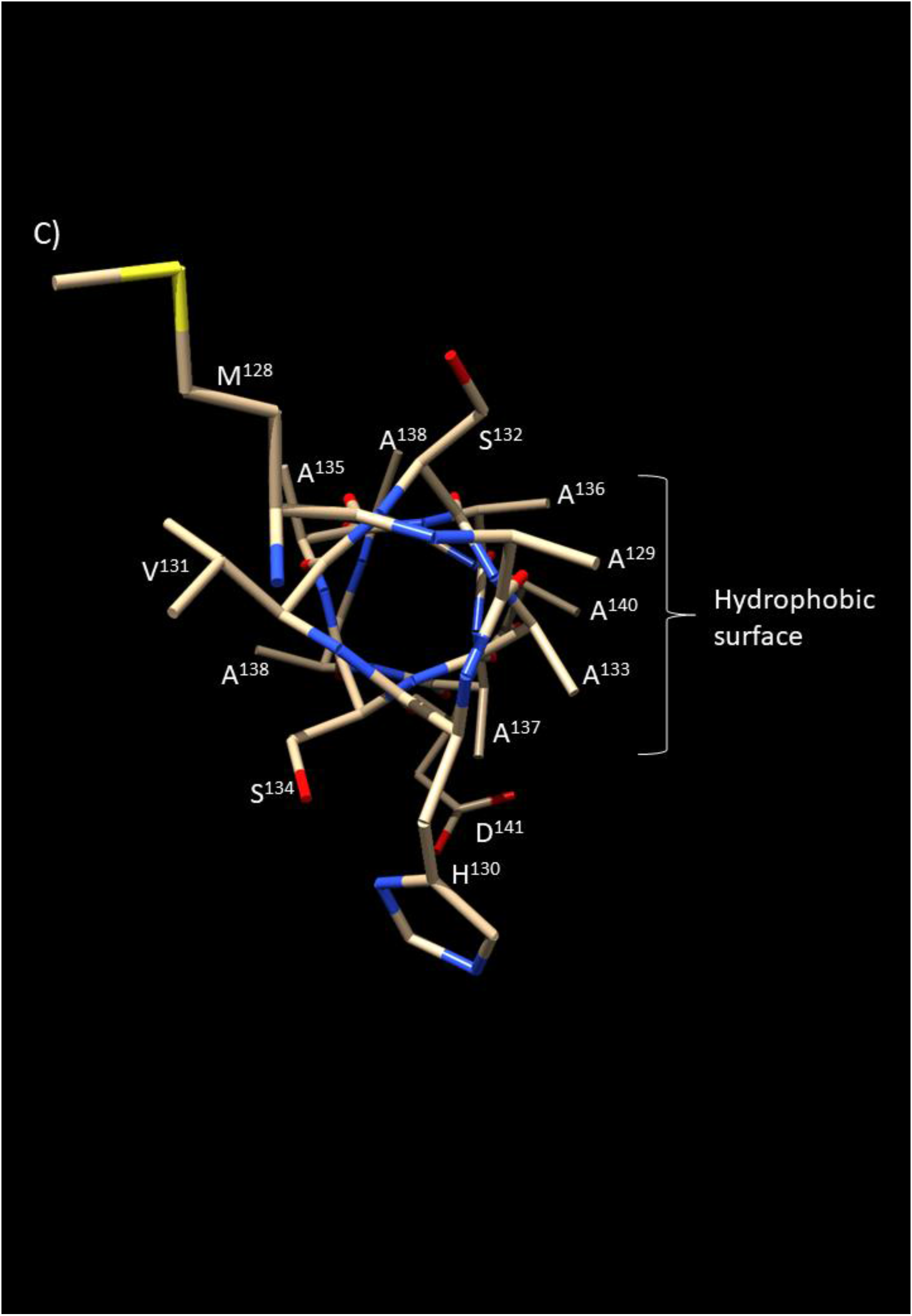
Structure of the 20-a.a. wild type IDR23 Ad12 peptide. The calculated structures for the wild type Ad12 peptide non-hydrogen atoms over the backbone and side chains (A) and Cα traces (B) are shown for residues using the single letter abbreviation and position N-(E^125^) to C-terminal (R^144^). Each figure represents the 14 lowest free energy conformers superposed from a conformational ensemble of 40 structures. In these final structures calculated, the average root mean square differences from the mean structure were 0.87 ± 0.12 Å for all atoms for the residues between M^128^ and D^141^ and 1.52 ± 0.17 Å for all the non-hydrogen atoms. No distance restraint was violated by more than 0.5 Å. (C) N-terminal end-on view of the backbone and non-hydrogen atoms within the average wild type Ad12 oncogenic spacer peptide structure from Met^128^ to Asp^141^ to highlight the amphipathic helix and hydrophobic surface presented by alanine residues at positions 129, 133, 136, 137 and 140.

### Effect of Mutation on the Structure of the Ad12 Spacer Peptide

It has been previously reported that substitution of a single alanine residue with proline (A^138^P) in Ad12E1A significantly reduces the oncogenic potential of a mutant virus designated *pm*715 (41, 43). To investigate this phenomenon, a theoretical model was derived for a 20-residue peptide carrying an equivalent Ala^14^→Pro substitution (Fig. 4A).

**Figure 4:**
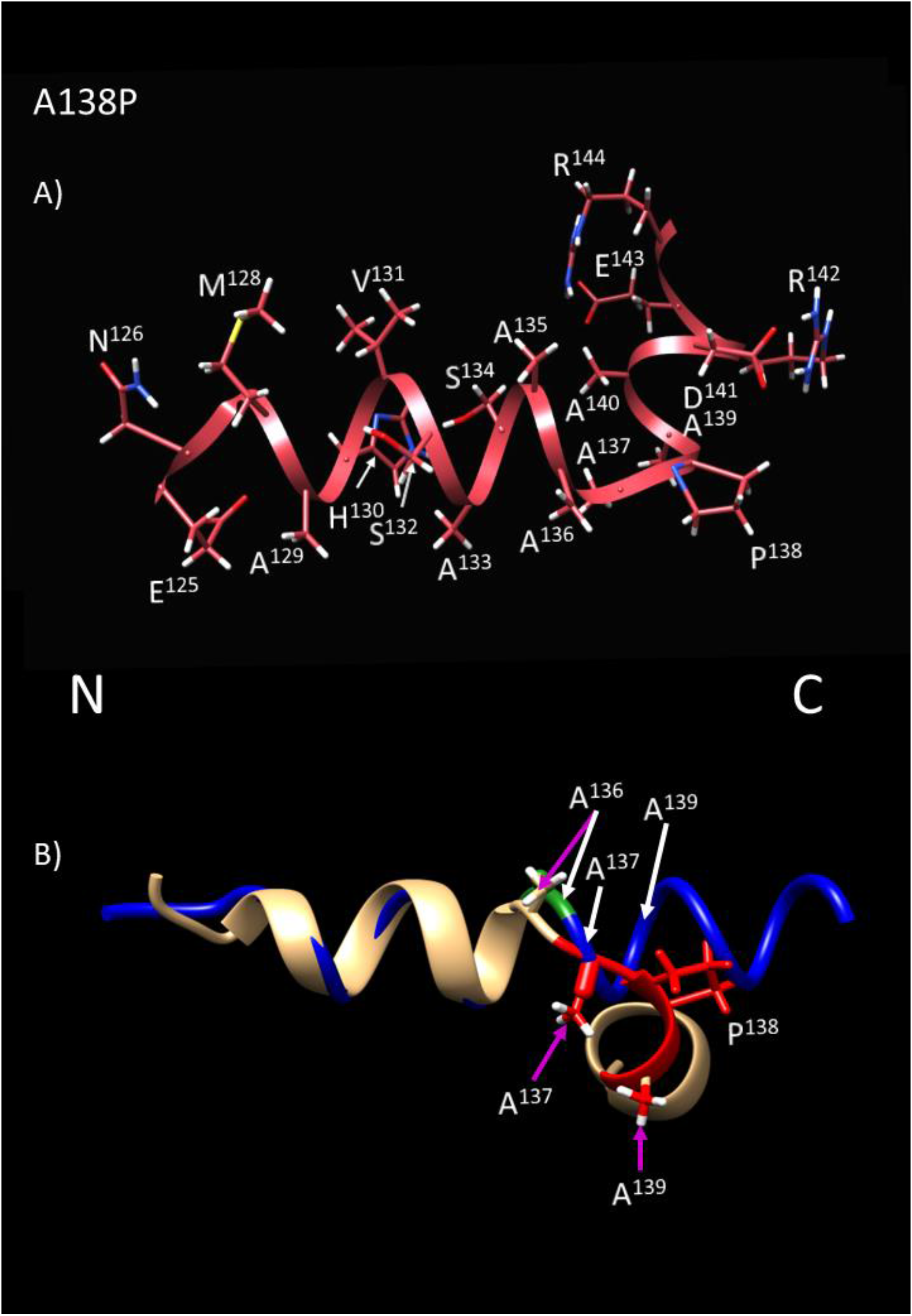
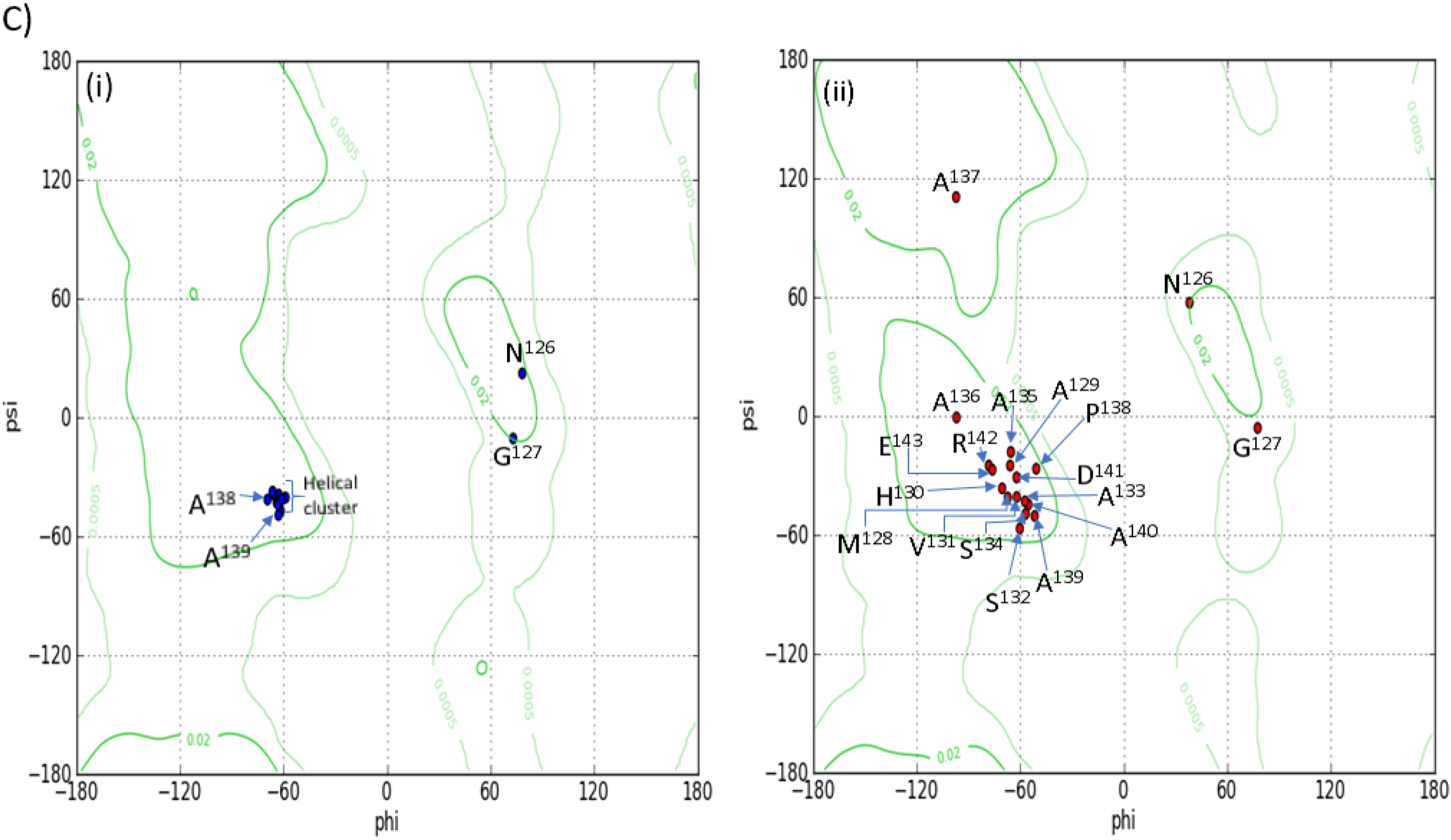
Theoretical structure of the 20-a.a. A138P isoform of the Ad12 peptide. A) The theoretical structure of a 20-residue peptide equivalent to amino acids E^125^-R^144^ of Ad12 E1A carrying the substitution of Ala^138^ to proline was generated using the I-TASSER theoretical modelling software. The backbone conformation is illustrated for residues between E^125^ (N terminus) and R^144^ (C terminus) as a red ribbon cartoon over the peptide backbone with amino acid side chains labelled according to type using the single letter abbreviation and position in the sequence of Ad12 13S E1A. B) The average structure calculated for the wild type Ad12 E1A-IDR23 peptide is presented as a Cα-trace (blue ribbon) superposed with the theoretical structure derived for the A^138^P form (gold ribbon) over the region E^125^(N)―R^144^(C). Amino acids are labelled as for (A). C) The Ramachadran plots for the average calculated structure for wild type (i) and theoretical A^138^P (ii) forms of the Ad12 E1A-IDR23 are presented.

The major structural consequences for the Ad12 wild type E1A-IDR23 peptide α-helical conformation when interposed with proline are predicted to include formation of a β-turn over residues A^135^―A^137^, N-terminal to the mutation and a significant juxtaposition of the C-terminal (Ala^139^―Asp^141^) α-helix perpendicular to the wild type peptide backbone axis (Fig. 4B). Conformational distortion within the P^138^ form of the peptide is also inferred from a wider dispersal of residues across the α-sector and notable translocation of Ala^137^ to the β-region in a Ramachandran plot (Fig. 4C). These predictions suggest that a proline residue introduced into E1A of the mutant virus is, in itself, disruptive of E1A-IDR23 three-dimensional structure and by inference, of the alanine-rich hydrophobic cluster that contributes to tumourigenesis. For these reasons, it was deemed that NMR spectroscopic data on the Ad12 P^138^ IDR23 peptide would be unlikely to provide much further insight.

Instead, the implications of a more conservative substitution, A^136^→S for Ad12 E1A-IDR23 structure were assessed, as a mutant Ad12 virus carrying an equivalent substitution is also defective in oncogenesis (41). Adopting a similar approach to that described above for wild type Ad12 E1A-IDR23, the structure of a 20-residue synthetic peptide carrying an equivalent Ala^12^→Ser substitution was determined (Fig. 5). Apart from A^11(135)^ and A^13(137)^, which flank the substitution, the chemical shift positions of backbone NH signals and NOEs observed were mostly equivalent between the two forms of the peptide (Fig. 5A). In this instance, structural integrity of the α-helix is maintained at position 12 of the peptide (labelled Ser^136^; Fig. 5 B and C). This finding supports the notion that although individual alanine residues may not necessarily be essential for maintenance of the helical nature of the Ad12E1A spacer region, integrity of the relatively hydrophobic surface formed by alanine residues is important for tumour development (Fig. 5D).

**Figure 5.**
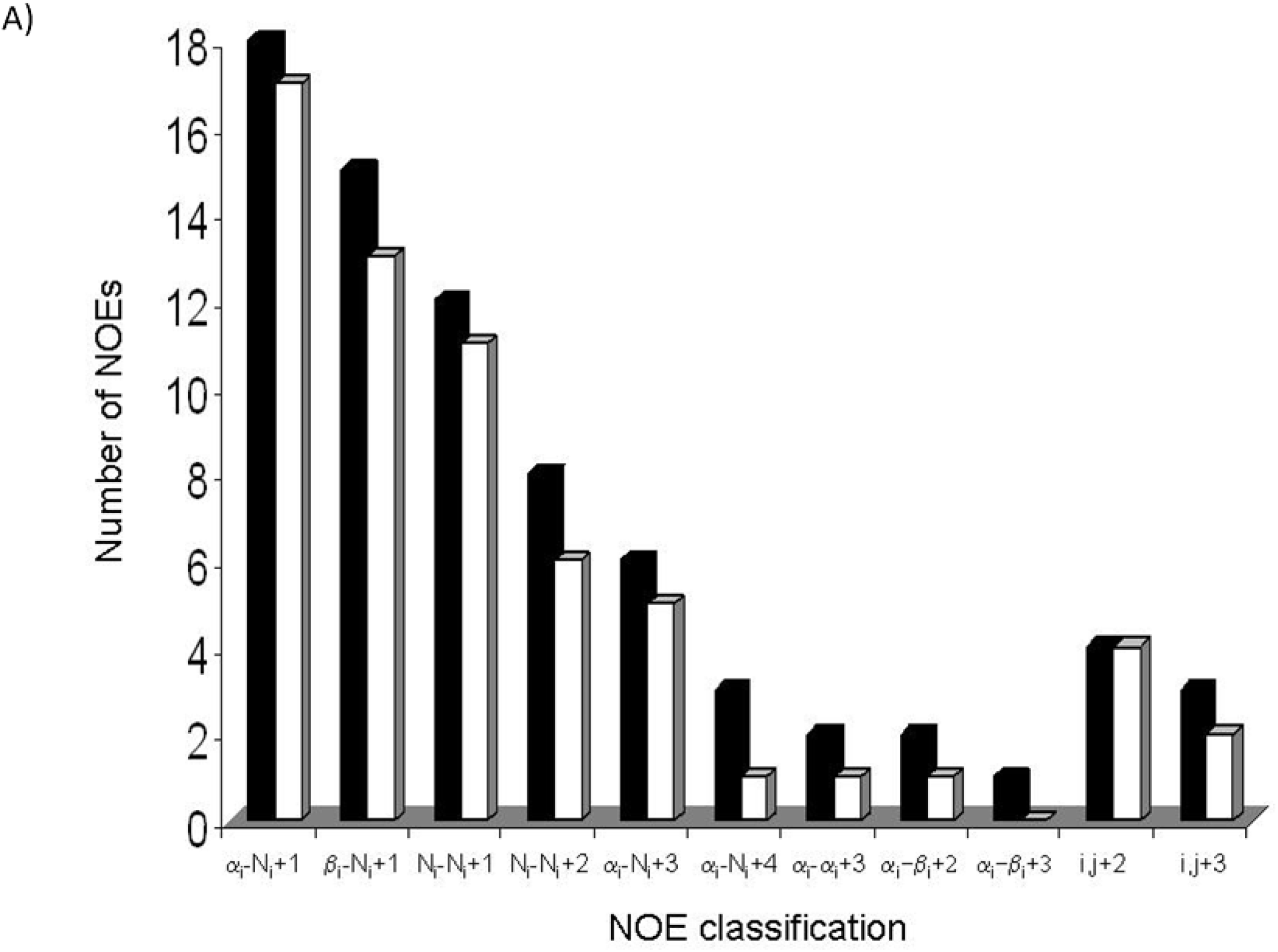
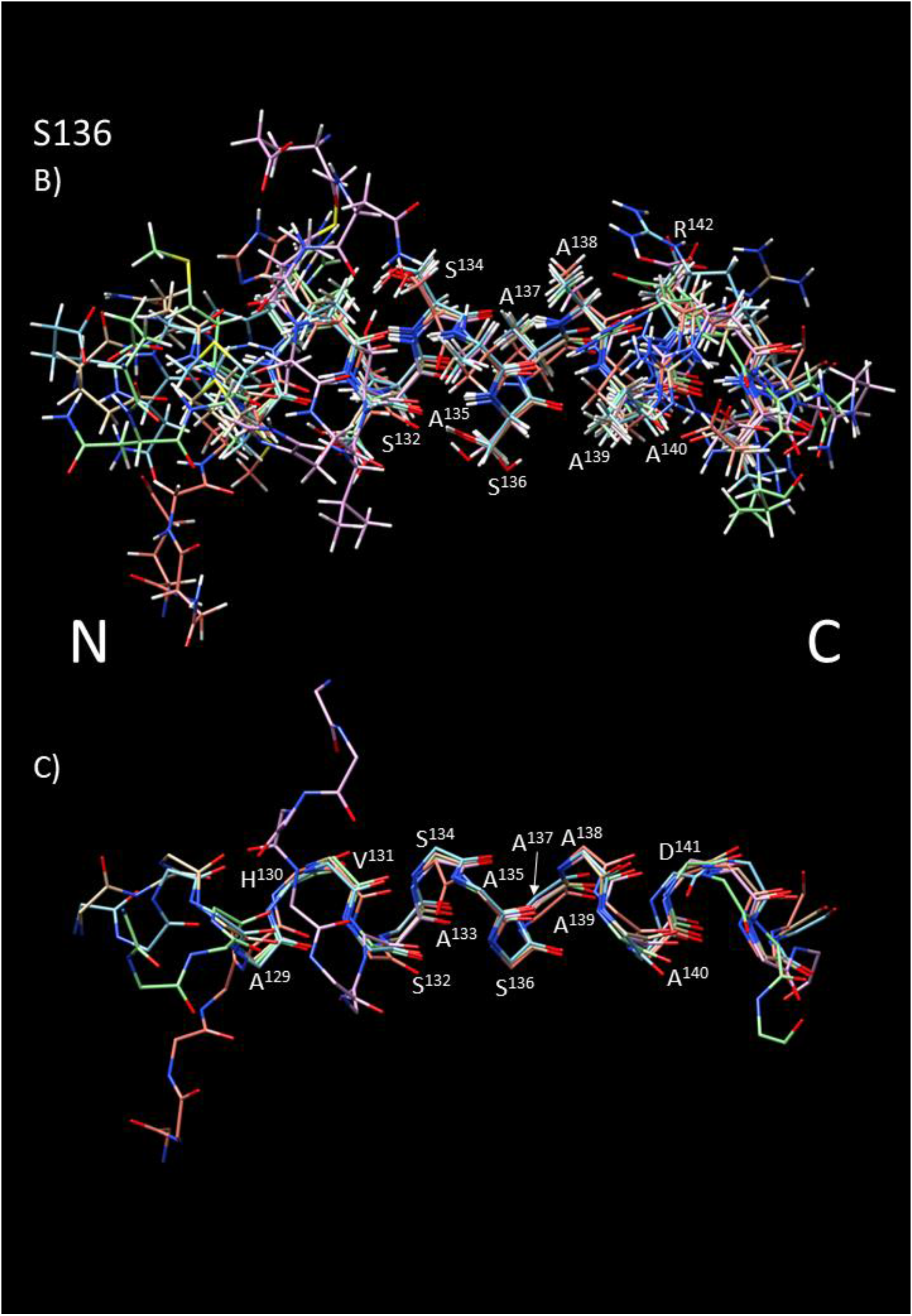
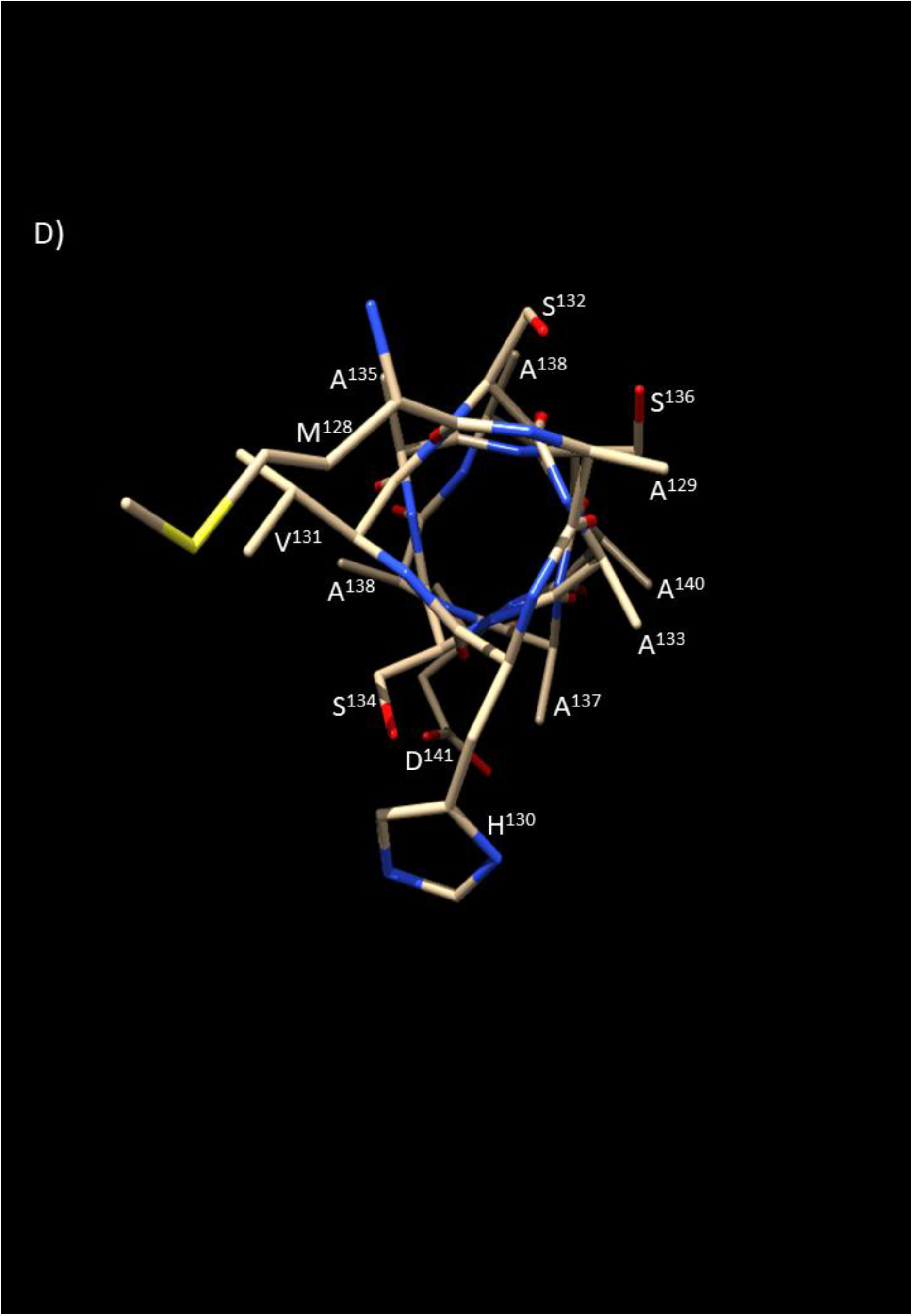
Structure of the 20-a.a. A136S isoform of the Ad12 E1A-IDR23 peptide. A) Summary of NOEs,^3^J _NHα_ coupling constants for the wild-type and A136S forms of the peptide. *wt,* black columns; mutant peptide, white columns. B) The calculated structure of S136 Ad12 E1A-IDR23. Nine superposed calculated structures for the A^136^S form of the Ad12 spacer peptide are shown as (A) non-hydrogen backbone atoms and (C) Cα-traces for residues between the N-terminus (E^125^) and C-terminus (R^144^). Amino acids are labelled using the single letter abbreviation and position in the sequence of Ad12 13S E1A. (D). Helical wheel representation of the Ser^136^ form of the Ad12E1A oncogenic spacer peptide used in this study. Amino acids are illustrated using the single letter abbreviation and position N-(E^125^) to C-terminal (R^144^) within the full length Ad12 13S E1A polypeptide.

### Sequence alignments of adenovirus E1As

Given the α-helical structure determined for IDR23 within Ad12E1A, it was of interest to consider the extent this might be conserved in E1A from other adenovirus species. A protein logo was generated from alignment of representative E1As from human and simian adenoviruses (reviewed; 29, 46) that herein emphasises a consensus motif of sequence ^9^M-A-X-A-S/A-X(2)-A/G-C/V-X-A-A/V-E^21^ (Fig. 6A). In addition, relatively low values for IUPRED disorder (61) in the range ≤0.58±0.01 were obtained for residues within various IDR23s equivalent to Ala^10^―Glu^21^ of the consensus (Fig. 6B). This finding could indicate that ordered structure is formed in the majority of E1A-IDR23s, consistent with that observed within the Ad12 E1A-IDR23 peptide (Fig. 3).

**Figure 6.**
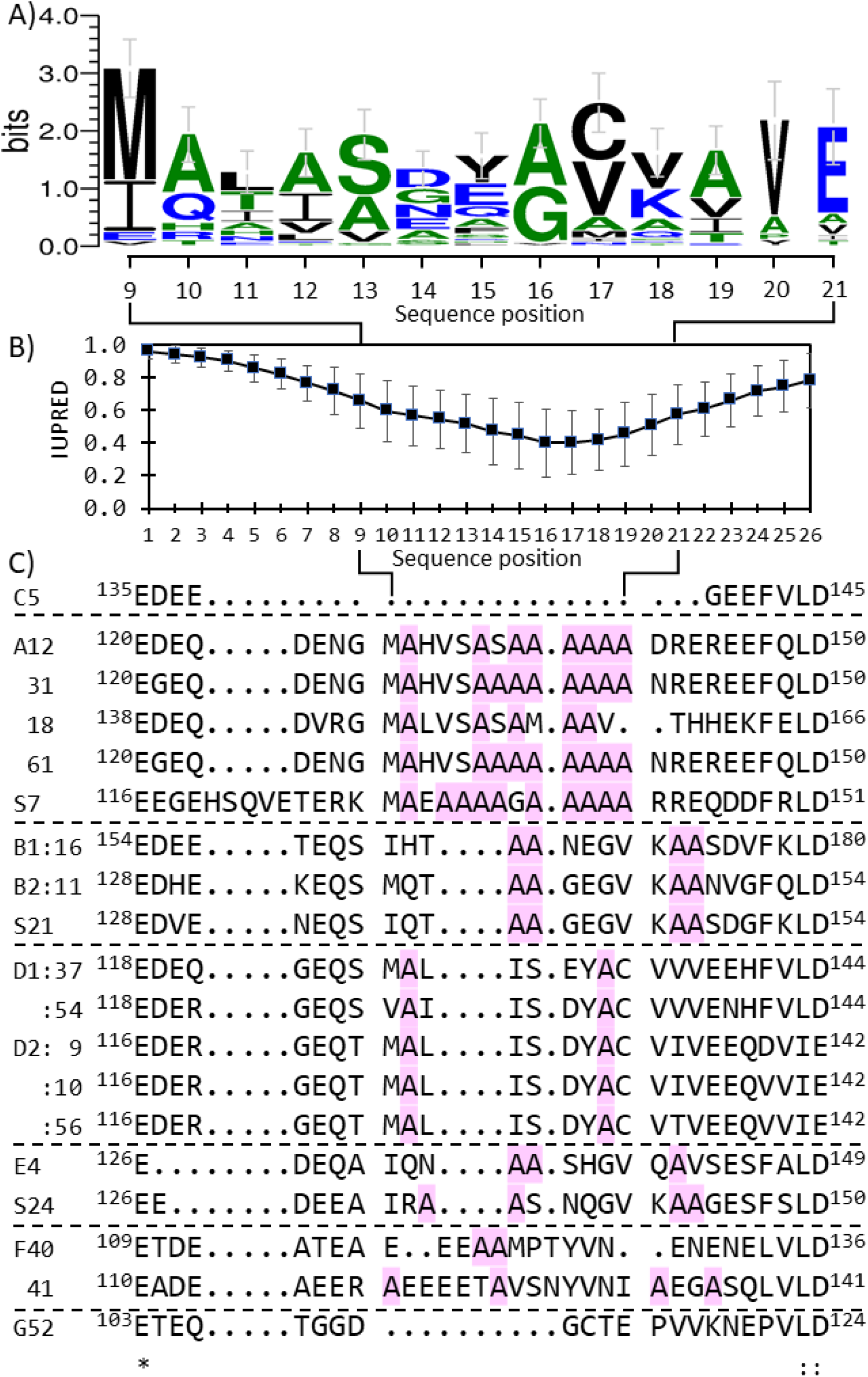
Amino acid sequence homology across AdE1A-IDR23 of representative viruses from subgroups A-G. A) The consensus sequence logo for 70 adenovirus E1A sequences is presented for each position between residues 9 and 21 in the alignment across IDR23 in units of bits with a maximum of 4.3 based upon the information content in the sequence of full length Ad5 13S E1A of 289 amino acids. B) Disorder predictions for individual amino acids in IDR23. Each datum point represents the average IUPRED score (±SD) for each amino acid within IDR23 from the N-terminus to the C terminus referenced to E^120^ and D^150^, respectively within Ad12 13S E1A and with score ranges between 0.0 for a fully ordered residue to 1.0 for a disordered conformation. C) Sequences of AdE1A-IDR23s shown are from just after the C-terminus of CR2 (E) to just before the N terminus of CR3 (L, D). The alignment is based on that for Ad12 E1A except that the Ad5 sequence has been transposed slightly. Viruses are from the following subgroups: Ad12, 18, 31, 61, SAdV-7; S7(A), SAdV-21 and 16(B1), 11(B2), 5(C), 37 and 54 (D1), 9, 10 and 56 (D2), 4 and SAdV-24 (E), 40 and 41(F) along with 52(G). The amino acid sequences from simian (S) adenovirus species 7i and 7ii which are homologous and designated S7, along with 21 and 24 shown are very similar to other members of their respective subgroups.

Specific details concerning the relative distribution of amino acids, notably alanine, across E1A-IDR23s were also obtained from the sequence alignments which here, commence at the C-terminus of CR2, Glu^135^ and finish at Asp^145^ within the N terminus of CR3 of Ad5 E1A (Glu^142^ of Group-D2 viruses; Fig. 6C). The alignment also includes slightly longer sequences from representative simian S7 and -F41 E1As, of 36a.a.’s and 32a.a.’s, respectively which, offset the overall configuration (Fig. 6C). With initial focus on the oncogenic Group-A viruses (Ad12, 18, 31 and 61), a motif of sequence ^9^M-A-X(2)-S-A-S/A-A-A/M-A(2)-A/V-A/X^21^ was observed (Fig. 6C). Ad12 and Ad31 E1A-IDR23s are closer in sequence identity than with Ad18 although, all three share alanine at positions 10, 14, 17, 18 and 19 in common with S7 E1A (Fig. 6C). These findings suggest that a hydrophobic surface formed by alanine residues arranged in an ordered helical conformation within IDR23 may be ubiquitous across oncogenic AdE1As.

E1A-IDR23 from the weakly-oncogenic SAdV-21 closely resembles counterpart sequences from subgroup -B1 and -B2 viruses, however, these in turn share only limited homology to the consensus motif with identity at Ala-12, −13, and −19 (Fig. 6C). Rather more radical substitutions occur in the replacement of ^17^A-A-A-A^20^ in Ad12E1A with ^17^C-V-V/I-V^20^ in Group-D1:54/-D2:9 E1As, respectively (Fig. 6C) which, would undoubtedly alter physicochemical/structural properties of the spacer region. Overall homology between E1A-IDR23s from non-oncogenic subgroups -E, -F and -G viruses is low and other amino acids within these are in the majority quite different, although SAdV-24 E1A aligns closely with HAdV-E4 E1A (Fig. 6C; 33). E1As from HAdV-F40 and -F41 appear to share alanine in common at positions 12, 13 and 19 and at 12 and 13, respectively to the consensus and HAdV-A12 E1A-IDR23 sequences (Fig. 6C). Alanine is absent within IDR23 of serotype-G52 E1A (Fig. 6C).

### Physicochemical properties of E1A IDR23s

Primary sequence alignments for AdE1A-IDR23s cannot be used exclusively to derive explanations for differences observed in adenovirus oncogenicity. These relate more to amino acid sequence variations that govern inherent protein folding propensities within the ‘spacer’ region and different solvent conditions in the cytosolic/nuclear environments within cancerous and normal cells (62). Thus, values of isoelectric points, pH(I) for E1A-IDR23 peptides were estimated using the ProtPARAM server (47). These are considered to range between 3.2 (lowest) and 4.4 (highest) for HAdV-F40 and -A18, respectively (Fig. 7A). These observations suggest that E1A ‘spacer’ regions are intrinsically acidic in general although, E1A-IDR23s from non-oncogenic viruses contain a higher percentage of glutamic acid residues and consequently, appear to be more acidic than oncogenic counterpart E1As (Fig. 7B).

**Figure 7.**
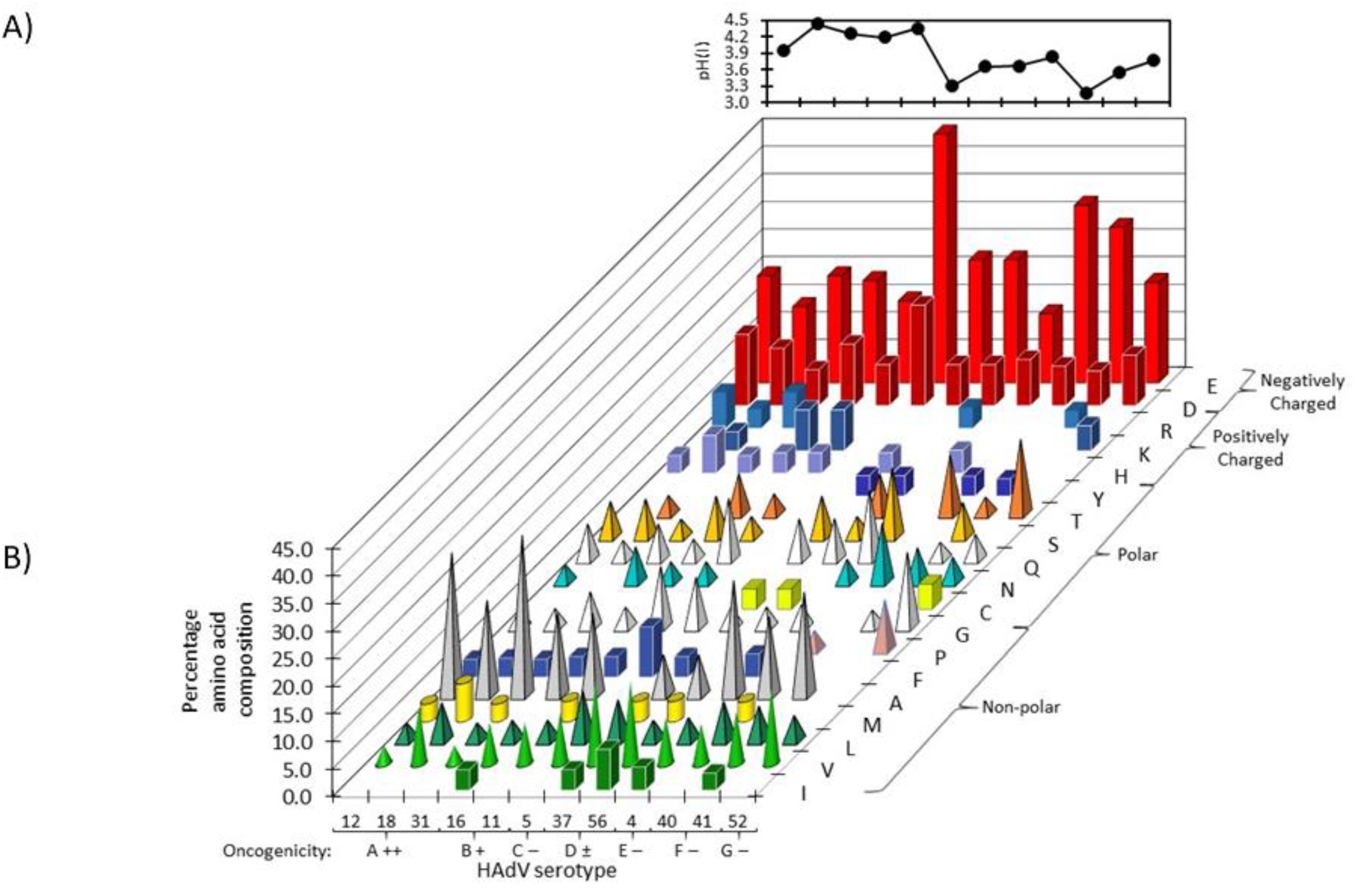
Physicochemical properties of amino acids within AdE1A-IDR23 from representative viruses from human subgroups A-G. A) The Isoelectric point pH(I) for E1A-IDR23 peptides from adenovirus subgroups A to G calculated using the ProtPARAM server are illustrated. B) The percentage amino acid composition for E1A-IDR23s from adenovirus species are presented according to polarity using the single letter abbreviation within highly oncogenic (++), weakly oncogenic (+ and ±), and non-oncogenic (-) human subgroups.

Inspection of the amino acid content within various E1A-IDR23s also revealed that alanine accounts for a higher percentage of total amino acids in the oncogenic Ad31 (29.03%) and Ad12 (25.81%) E1As, compared to either Ad18, the weakly-oncogenic -B1 and -B2 and non-oncogenic Group-D AdE1As studied (17.24%, 14.81%, and 7.41%, respectively; Fig. 7B). Interestingly, the hydrophobic amino acid, isoleucine is absent from spacer regions in the majority of oncogenic Group-A/B and non-oncogenic -F40 and -G52 AdE1As (Fig. 7B) whilst positively charged lysine and arginine residues are absent within E4 E1A-IDR23 which, therefore, appears to be strongly acidic (pI = 3.83; Fig. 7A). Although alanine accounts for 20.83%, 14.29% and 18.75% of total amino acids in E1A-IDR23 from –E4, -F40 and -F41 viruses, respectively (Fig. 7B) these are of comparatively shorter sequence length compared to the oncogenic counterparts within the relative alignment (Fig. 6C).

### Hydrophobicity within IDR23 of E1A from different adenoviruses

Further physicochemical properties of amino acids within IDR23 of various E1As were obtained from calculations of hydropathy or GRAVY score (63). This parameter represents a quantification of hydrophobicity, or polarity for all amino acids as a function of polypeptide sequence length (63). Overall, E1A-IDR23s from Group-C and oncogenic Group-A/B viruses appear to be hydrophilic whilst, non-oncogenic Group-D counterparts appear to be comparatively hydrophobic (Fig. 8). Specifically, a GRAVY score of −1.05 was calculated for Ad12 E1A-IDR23 which, appears to be acidic/hydrophilic, Ad18 E1A significantly less so (−0.34) and Ad31 E1A theoretically aligns with weakly-oncogenic Group-B virus E1As which appear to be slightly more hydrophobic (*ca*~ −0.8; Fig. 8). Exceptionally, E1A-IDR23 from the oncogenic -D2:9 adenovirus appears to be less hydrophobic than other non-oncogenic Group-D species and aligns more closely with oncogenic Group-A/B E1A-IDR23 peptides (Fig. 8). These findings arise as a consequence of differences in the ratio of hydrophobic (Ile and Val *cf* Ala) to charged (Lys/Arg and Asp/Glu) amino acids between E1As and potentially indicate that variations in structural properties exist between different E1A-IDR23 peptides. Hydrophobicity scores were also calculated for spacer region of E1As from the non-oncogenic adenoviruses -F40, -F41, and -G52 and found to be −1.03, −0.69 and −0.89, respectively which are theoretically equivalent in hydrophobicity to the hydrophilic Group-A/B AdE1As (Fig. 8). Nothwithstanding, IDR23 from E1A of the non-oncogenic HAdV-E4 serotype is of similar polarity to the group D viruses (Fig. 8) although, the sequences are quite different (Fig. 6C).

**Figure 8.**
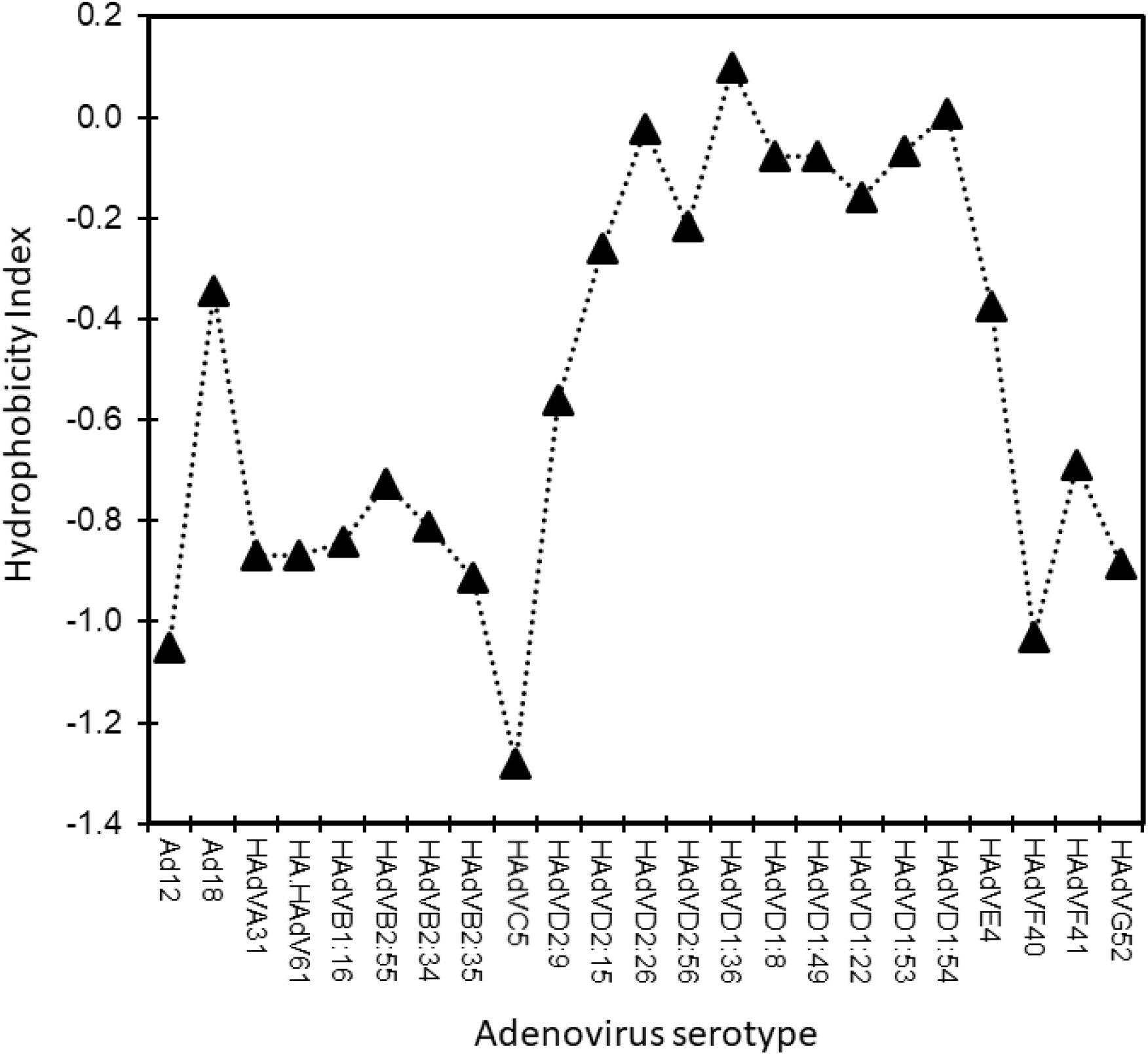
Hydropathy index for AdE1A-IDR23s from representative viruses from human subgroups A-G. The grand average for hydropathy or GRAVY score calculated for individual E1A-IDR23 spacer regions from human adenovirus species using the ProtPARAM are presented. The scores range between negative values for hydrophilic and positive for more hydrophobic peptides.

### Structure predictions of IDR23s from different adenovirus serotypes

Further analyses of IDR23s from various E1As involved derivation of a protein logo and predictions for of β strand (E), helix (H) and random coil (C) secondary structural elements according to adenovirus subgroup (Fig. 9). For the non-oncogenic consensus Ad5E1A sequence where IDR23 is essentially absent, a short α—helix was predicted for residues ^135^EGEE^141^ in between CR2 and CR3 (data not shown). As expected, predictions of secondary structure vary between representative E1As of the other serotypes investigated.

**Figure 9.**
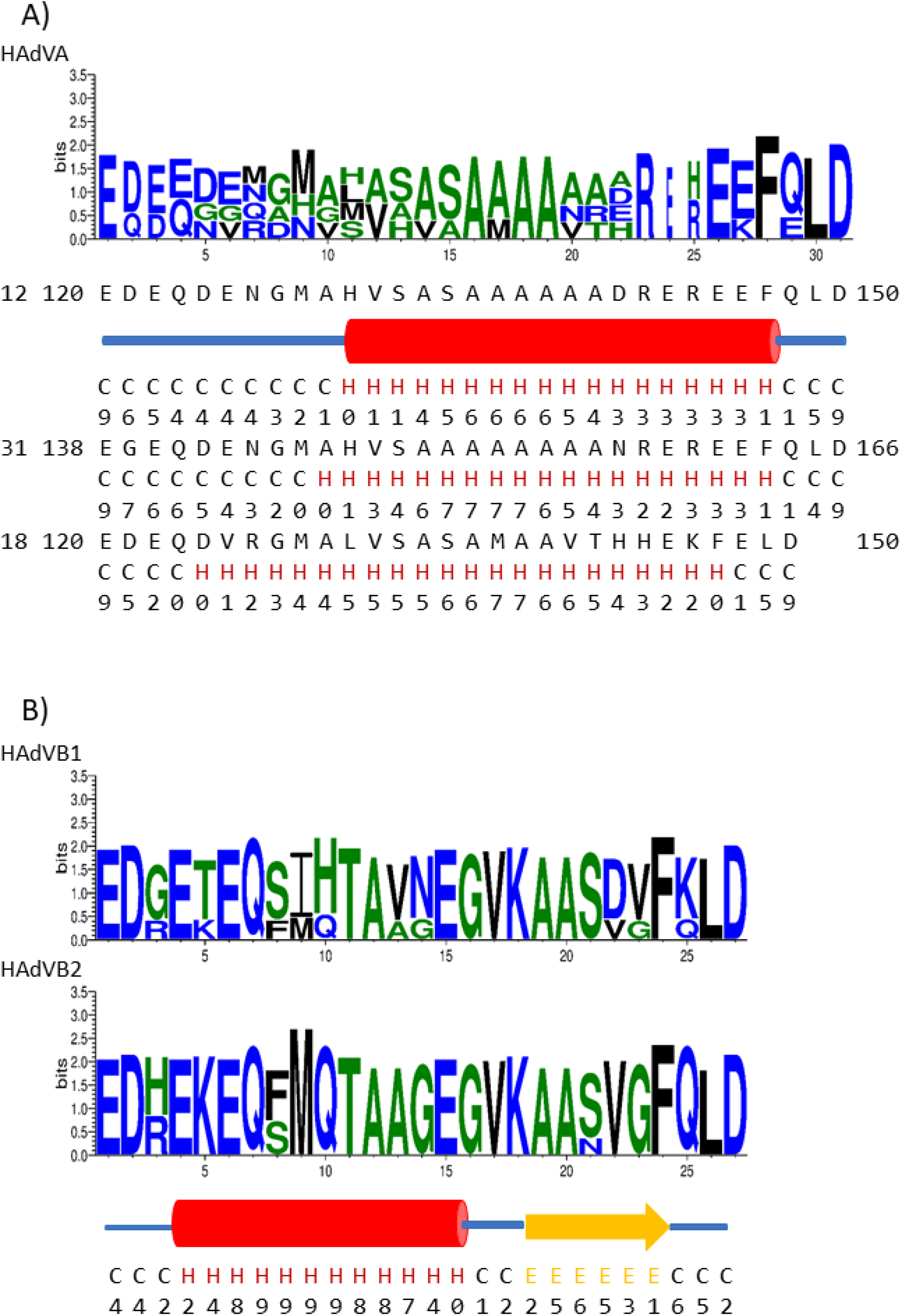
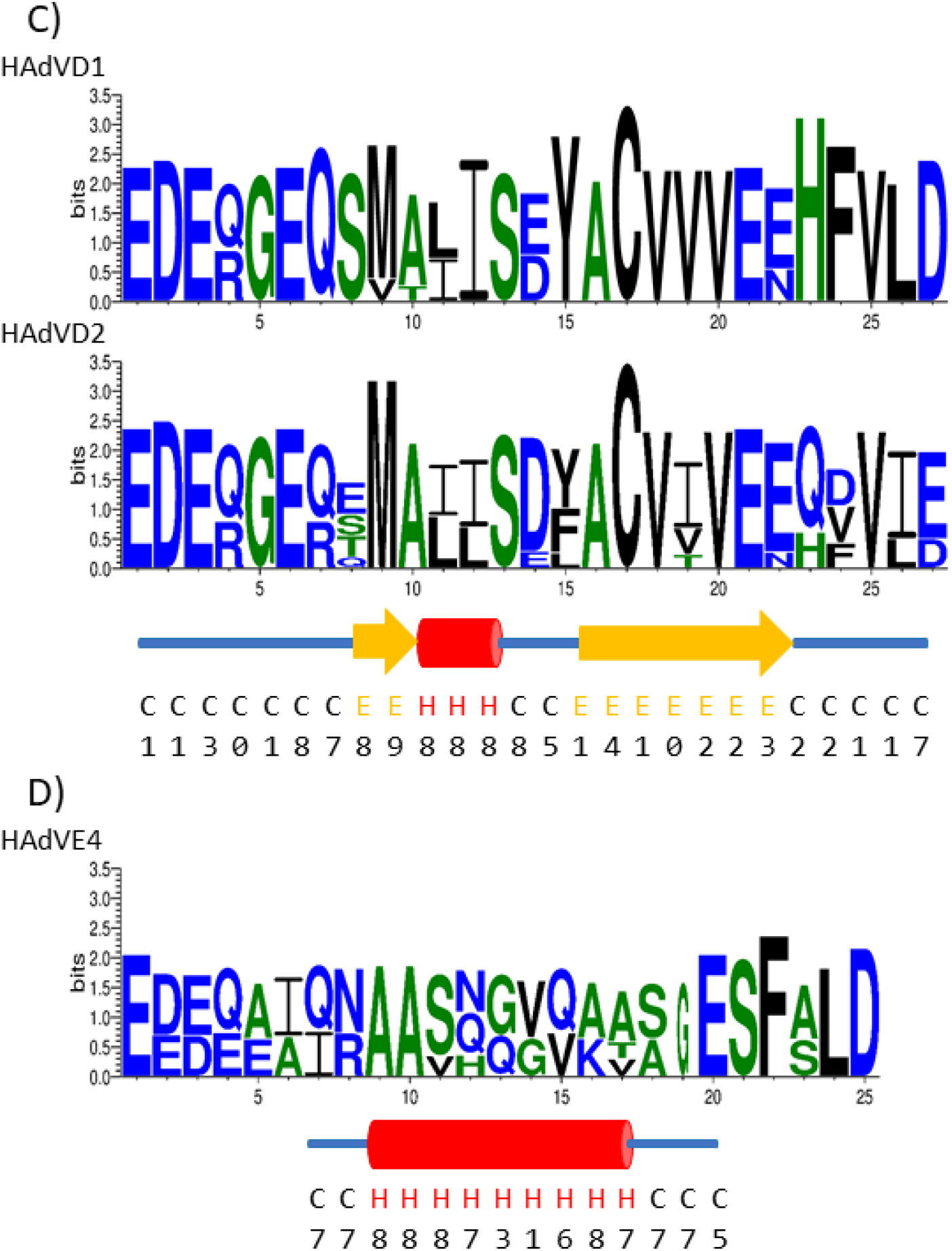
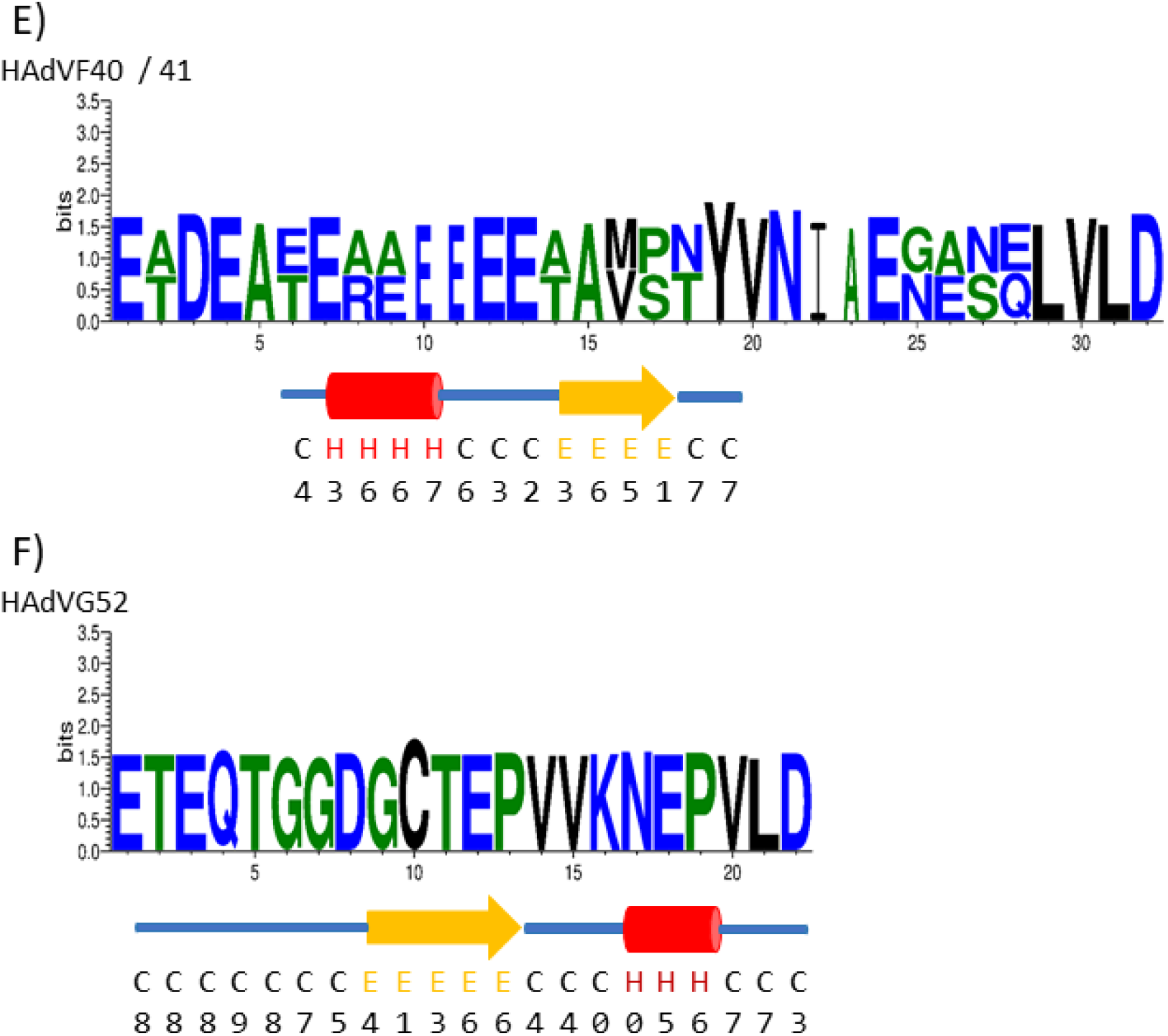
Predicted conformation of the IDR23 spacer regions from representative human adenoviruses from subgroups A-G. Sequence logos for E1A spacer regions according to adenovirus serotype sequence alignments are shown together with the predicted structures (H = helix, E = β sheet/turns and C = random coil). Degree of confidence of the prediction is also shown over each sequence. Red cylinder shape, helix; broad yellow arrow, β turn and blue line, coil. Data are shown for (A) AdE1As from subgroup-A12, 31 and 18; (B), subgroups B1:7 and 16 and B2:34, 35 and 55, (C) sub group D1:8,22,36 and 49; D2:9, 10, 15; (D) subgroup-E4; (E) subgroup-F40 and 41 and (F) subgroup-G52.

Examination of the oncogenic HAdV-A serotype E1As highlights the presence of an alanine-rich sequence within the sequence ^9^M⎯A/D/E/H^22^ of the protein logo and an α-helix is predicted with high confidence to extend over residues H^130^⎯F^147^ of Ad12 E1A (Fig. 9A), consistent with the NMR data (Fig. 3) and previous studies (56). When all other group-A viruses were examined, several differences to Ad12 were noted (Fig. 9A). In Ad18E1A, a more extensive α-helix is predicted to span the sequence equivalent to D^124^ (at the C terminus of CR2) to F^147^. In contrast, the helix is less extensive in Ad31E1A and more similar to Ad12E1A over the region A^147^―F^163^. In all three cases, helices are bounded by regions of random coil and the helical extent does not correlate well to the proportion of alanine residues (Fig. 9A).

In consideration of E1A spacer regions from adenovirus subgroups-B1 and -B2, identity between these and group-A E1A counterparts is low (Fig. 9B). The N-terminal half from each is predicted to be α-helical over residues ^4^Glu⎯Glu^15^ followed by a short span of random coil and a β-strand that extends over the sequence ^19^A-A-S-D/V-V/G-F^24^ (Fig. 9B). Within representative E1As from HAdV-D1 and -D2, a short β-strand is predicted over the sequence ^8^M―A^10^ followed by a helix between ^11^L/I and S^13^ and a β-strand over ^16^A―E^21^ of sequence logo (Fig. 9C). Interestingly, an α-helix is predicted within IDR23 of Ad4E1A although, there is very little sequence conservation to the Group-A viruses (Fig. 9D). In Ad40 and Ad41 Group-F serotypes, a rather less extensive α-helix is predicted over the N-terminus of the sequence motif, ^7^Glu-Ala-Ala-Glu^10^ separated by a short extended region from a β-strand over the C-terminal region ^14^A/T―P/S^17^ (Fig. 9E). Within IDR23 from the non-oncogenic Ad-G52 E1A, which shares no identity with the consensus sequence, secondary structural elements over the regions ^9^G⎯P^13^ and ^16^N⎯P^19^were predicted to be a β-strand and an α-helical conformation, respectively (Fig. 9F).

## Discussion

Soon after the first scientific and historically significant demonstration that a human adenovirus could cause tumours, albeit in rodents, studies focused on Ad12 and Ad18 serotypes (3–5). Later on Ad7, was also observed to cause tumours in newborn rodents but at a lower frequency and with a much longer latency period (8). As more species of adenovirus were isolated, it became evident that members of the Group-A viruses are oncogenic, Group-B viruses are weakly-oncogenic and the great majority, most notably Ad5, are non-oncogenic in immunocompetent animals.

The most obvious primary structural difference between Ad5 and Ad12 E1As is the presence in the latter protein of a short spacer sequence, rich in alanine residues between conserved regions 2 and 3 (32, 33) designated IDR23 (29). Deletion of this sequence from Ad12E1A causes a marked reduction in the ability of the resulting mutant virus to cause tumours in newborn rodents (41, 43). However, it was noted that this virus retained a very limited tumour-inducing activity and this led to the notion that other sequences, most probably the N-domain of Ad12E1A, also contributes a certain extent to the oncogenic potential of the virus (43). Significantly, it has also been shown that the Ad12E1A spacer region is not responsible for the Ad12E1A-mediated down regulation of MHCI on the surface of Ad12 transformed cells, which had previously been considered to be of considerable significance in determining the ability of Ad12 to induce tumours (16, 22, 64).

In an attempt to understand the role of IDR23 and, in particular, the essential differences between the equivalent regions in AdE1As from non-, weakly- and oncogenic subgroups, the present study was undertaken. In structural predictions of AdE1As from several subgroups, it was reported that all spacer regions are likely to be unstructured (28). However, in a more limited study we suggested that the Ad12 spacer is very likely to be, at least partially, helical (56), consistent with low values of intrinsic disorder calculated for amino acids in IDR23 reported here and elsewhere (29). In an attempt to resolve this contradiction, using NMR spectroscopy, we have determined the structure of a synthetic peptide identical to the Ad12 spacer region. This was found to be predominantly α-helical (Fig. 3) as might be expected for a sequence very rich in alanine residues (65). In this present study, the 3-D structure of a peptide equivalent to IDR23 from Ad12 E1A was determined using trifluoroethanol as a co-solvent with water, since it is well known that this solvent can enhance solution structures by promoting hydrogen bond formation within the peptide (66). TFE also reduces the formation of spurious secondary structures through hydrophobic aggregation (67). It is also noteworthy that our observations were, on the whole, similar for NMR spectral data for the Serine 136 form of the peptide (Fig. 5). Here, introduction of a serine residue within the poly-alanine sequence leads to only a very limited disruption of the amphipathic surface nature of the α-helix around the “mutation”, rather than disruption of the intrinsic helical structure itself (Figs. 3 and 5).

Structural predictions for a peptide containing a substitution (A^138^→P) identical to the mutant virus, *pm*715 which, has a reduced ability to cause tumours in rodents (41, 43) indicated that there would be a marked disruption of helical propensity around that substitution and over the C-terminus of the peptide (Fig. 4). By convention, the N―Cα (φ) dihedral torsion of proline consigns this amino acid to a ‘helix-breaker’ function (68) and this alone, is responsible for loss of tumour-inducing activity as presumably, Ala^138^ directly participates in a protein•protein interaction essential for tumourigenesis.

Comparison of the predicted structures of the spacer regions of three subgroup-A viruses shows variation of helical content with Ad31 being closer to Ad12 than with Ad18 (Fig. 9). The helical content within the Ad18 peptide is predicted to be more extensive although, this serotype is reported to be rather less tumourigenic than Ad12 (8). An increase in α-helical content predicted within IDR23 of Ad18 E1A most likely occurs as a consequence of increases in the content of histidine, leucine, methionine and valine at the expense of alanine. The fact that HAdV-A12 expressing either Ser^136^ or Pro^138^ E1A display appreciably reduced levels of oncogenicity (41, 43) indicate that integrity of the hydrophobic surface formed by alanine residues in a helical conformation is of central importance to that property.

It is noteworthy that E1A-IDR23 peptides vary significantly in sequence length and amino acid composition (Fig. 7). Indeed, variations in the ratios of IMLV to DE/RK residues within different E1As may be of particular influence where the weakly hydrophobic amino acid alanine is clustered on the hydrophobic side of an amphipathic helix (65). For example, the cumulative hydrophobicity of +9 for alanine residues converged on one side of an amphipathic helix within Ad12E1A-IDR23 would marginally increase to 9.7 with the substitutions Ala^17^→Met and Ala^20^→Val in Ad18E1A-IDR23 (63). Any disruption to target protein binding interactions would most likely arise as a consequence of steric hindrance associated with the large side chains of Met and Val. Furthermore, replacement with Ser^136^ would only marginally reduce the amphipathic helix apolar surface hydrophobicity by −0.8 (63). The mere presence of the Ser―^β^OH moiety is also indicative of surface area restrictions imposed by alanine(s) on target protein associations. Although, the molecular and/or signalling events of these required for tumour formation will have to await further investigation.

IDR23 from Group-B virus E1As are predicted to contain a more limited N-terminal region of α-helix that precedes a β-strand (Fig. 9B). It has previously been demonstrated that Ad7 (a Group-B1 virus) could cause tumours in newborn hamsters but only after a considerable incubation time (about 3-4 times as long as for Ad12; 8). From a consideration of spacer regions of the Group-B (weakly-oncogenic) viruses, it can be seen that a limited number of alanine residues are conserved. In the subroup-B2 viruses there are 4 alanine residues in the spacer whereas there are 3 in the -B1 viruses (Fig. 7C). However, it remains unclear, on the basis of these results, whether maintenance of the α-helical nature of E1A-IDR23, or very limited sequence homology is the limiting factor for a lower frequency tumours seen with Group-B viruses in general (8, 9).

From the structural predictions presented (Fig. 9), it can be seen that amino acids within representative E1A-IDR23s from Group-D viruses are predicted to form both helical and β-conformational structures. It can, therefore, be concluded that the simple presence of an extended region of α-helix in the spacer is not sufficient to render the protein oncogenic. Therefore, position and orientation of particular amino acid side chains must be a determining factor for this although there is probably also a requirement for an α-helical structure in the spacer that enables efficient tumourigenesis. Within unaligned sequences of, for example, Ads –A12 and -E4, two alanines at positions 10 and 16 are equivalent, although considerable variation over the hydrophobic surface of an amphipathic helix would be expected (data not shown). Again, the observation that substitution of an alanine for a proline in *pm*715 E1A that results in a significant reduction in tumour forming ability (41) and major disruption to the apolar surface formed on one side of the helix supports this contention (Fig. 4).

It seems likely that the Ad12E1A spacer region is a site of interaction with various unknown cellular proteins, although it is possible that the integrity/structure of IDR23 is prerequisite for correct orientation and the binding to novel cellular targets. Significantly it has been shown, using microarray analysis of Ad12 transformed rodent cells, that the intact AdE1A spacer region is responsible for increased expression of neuronal-specific genes that are associated with cell cycle regulation, transcription regulation and tumour invasion (69). The mechanism for this remains unclear although, it could plausibly involve interactions of the spacer with one or more, as yet unknown, cellular regulators. Similarly, differences in structural propensity of E1A-IDR23s and target protein interactions will have to await further investigation.

It might be expected that the amino acid sequence of the spacer region would mirror sequences found in certain cellular proteins in the same way that short sequences in other regions of E1A duplicate cellular protein binding sites, for example, the Rb family binding site in CR1 (30) and the CtBP binding site in CR4 (55). Poly-alanine sequences are common in mammalian proteins, occurring in, for example, a number of homeobox and zinc finger proteins as well as various transcription factors (70). If the IDR23 sequence logo from the Group A E1As ^9^M-A-X(2)-S-A-S/A-A-A/M-A(2)-A/V-A/X^21^ (Fig. 7C) is used to scan protein databases, a number of proteins containing identical motifs can be identified. These include homeobox proteins (e.g. HMX3, VAX1, and HOXA13) and RNA binding proteins, such as RBM24, as well as a variety of other cellular components. No proteins resembling the Ad7 and Ad11 (Group-B) spacer motifs were found (data not shown). In the case of SAdV-7, a poly-alanine sequence with a single glycine residue shares homology to a few proteins including, histone-lysine N-methyl transferase 2. The similarity of the Ad12 spacer region to proteins involved in transcriptional regulation is consistent with the observations that there are marked changes in protein expression in response to the spacer region (28). It is quite possible that the interaction of one or more of the potential E1A homologues is disrupted in the presence of the Group-A IDR23 motifs and a novel deleterious association formed. Unfortunately, such experiments are limited with high uncertainty in identifications for target proteins, particularly when only a very small number of amino acids are needed to be identical.

In summary, we have demonstrated that a peptide identical to the Ad12 spacer region is α-helical. Substitution of one of the alanine residues causes disruption of the helix, its degree depending on the residue introduced. It is predicted that spacer regions from other subgroups have limited helical content and it has been concluded that this conformation is necessary for high levels of oncogenicity although there appears to be some requirement for specific amino acid side chains.

## Acknowledgements

We are most grateful to Chongqing Medical University and Cancer Research UK for financial assistance.

## Abbreviations

Ad: adenovirus
CR: conserved region
E1: early region 1
E1A-IDR23: inter domain region 2-3 or early region 1A protein oncogenic spacer
Hz: Hertz
NMR: nuclear magnetic resonance
NOE: nuclear Overhauser effect
NOESY: nuclear Overhauser effect spectroscopy
TBP: TATA binding protein
TFE: trifluoroethanol
TOCSY: total correlation spectroscopy

## Table One: Adenovirus serotypes

The early region 1A proteins used in this are listed according to species, serotype and relative oncogenicity.

